# PPFlow: Target-Aware Peptide Design with Torsional Flow Matching

**DOI:** 10.1101/2024.03.07.583831

**Authors:** Haitao Lin, Odin Zhang, Huifeng Zhao, Dejun Jiang, Lirong Wu, Zicheng Liu, Yufei Huang, Stan Z. Li

## Abstract

Therapeutic peptides have proven to have great pharmaceutical value and potential in recent decades. However, methods of AI-assisted peptide drug discovery are not fully explored. To fill the gap, we propose a target-aware peptide design method called PPFlow, based on conditional flow matching on torus manifolds, to model the internal geometries of torsion angles for the peptide structure design. Besides, we establish a protein-peptide binding dataset named `PPBench2024` to fill the void of massive data for the task of structure-based peptide drug design and to allow the training of deep learning methods. Extensive experiments show that PPFlow reaches state-of-the-art performance in tasks of peptide drug generation and optimization in comparison with baseline models, and can be generalized to other tasks including docking and side-chain packing.

## 1. Introduction

Therapeutic peptides are a unique class of pharmaceutical agents composed of a series of well-ordered amino acids. The development of peptide drug design and discovery is accelerated by fast advances in structural biology, recombinant biologics, and synthetic and analytic technologies (Wang et al., 2022). Peptide drugs play an important role in pharmacology because they usually bind to cell surface receptors and trigger intracellular effects with high affinity and specificity, showing less immunogenicity and taking lower production costs (Muttenthaler et al., 2021). Therefore, they have raised great research interests as new peptide therapeutics are continuously developed with more than 150 peptides in clinical tests and another 400–600 peptides undergoing preclinical studies.

Deep learning has revolutionized fields like drug discovery and protein design (Makhatadze, 2021; Watson et al., 2023; Dauparas et al., 2022), which proves to be effective tools to assist the development of small and large molecule drugs. Peptide drugs occupy a unique chemical and pharmacological space between small and large molecules, but the AI-assisted peptide drug discovery methods remain limited compared with those established for small molecules and proteins. Unbound peptide chains are usually at high free energy and entropy values, thus showing unstable conformations, while they trigger pharmacological effects when binding to specific receptors, forming a complex with equilibrium structures that are composed of a pair of receptor and ligand. (Marullo et al., 2013; Seebach et al., 2006). Therefore, we focus on the designation of peptide drugs that can bind to specific receptors (Todaro et al., 2023). Recently, structure-based drug design (SBDD) methods are developed for target-aware small molecule generation (Peng et al., 2022a; Guan et al., 2023; Lin et al., 2022), while these methods cannot be simply transferred to peptide design tasks for the following reasons: (i) Peptide drugs are usually of larger molecular weights (500 − 5000 Da) compared with small molecule drugs (< 1000 Da) (Francoeur et al., 2020); (ii) Topologies of atom connectivity in peptides are close to proteins rather than molecules; (iii) Internal geometries in peptides are of different patterns from small molecules.

Hence, we identify four challenges for the target-aware peptide drug design. *First*, how to extract sufficient information from the conditional receptor contexts. This is indispensable for the model to generalize to new receptors. *Second*, how to generate valid peptides that satisfy physicochemical rules. If the generated peptides are not chemically valid, other drug properties are unnecessary for further consideration. *Third*, searching for natural peptides and replacing them with animal homologs, such as the discovery of insulin, GLP-1 and somatostatin, were the important strategies used for peptide drug discovery (Kelly et al., 2022). Therefore, instead of de novo design, the model should apply to another scenario: optimizing natural peptides to drugs with higher binding affinities. *Besides*, there are few available benchmark datasets large enough to support the training of deep learning models for structure-based peptide drug design tasks, and it is urged to collect and organize a high-quality dataset that satisfies the demand for massive data.

(*i*) *To address data shortage*, we first construct a dataset consisting of a large number of protein-peptide complexes through a series of systematic steps. (*ii*) *To give a solution to the task*, we propose a generative model based on flow matching, to learn the structure distribution of peptides from their torsion geometry and the distributions of global translation and orientation. Instead of directly learning the explicit probability, PPFlow fits the gradient fields of the variables’ evolutionary process from a prior distribution to the peptides’ sequence-structure distributions and uses the learned gradient field to iteratively push the sequences and structures toward data distribution by an ordinary differential equation. (*iii*) *To sufficiently extract representation from the protein*, we employ neural networks as the approximation to the gradient fields, parameterized with attention-based architectures to encode contextual proteins’ structure and sequence. (*iv*) *To confirm the validity of the model*, extensive experiments on multi-tasks are conducted. PPFlow reaches state-of-the-art performance compared with diffusion-based methods we transferred in generating peptides with higher affinity, stability, validity, and novelty. In optimization tasks, PPFlow also shows great superiority. Besides, experiments on flexible re-docking and side-chain packing demonstrate the models’ potential in structure modeling.

To sum up, our contributions include: (i) **New Task**. We consider the conditional generation of peptide drugs targeted at protein receptors as contexts, which is an area that has rarely been explored by deep learning. Other tasks including peptide optimization, flexible re-docking, and side-chain packing are also studied and evaluated. (ii) **Novel Model**. To fulfill the task, we establish a new model called PPFlow as a complete solution to model the internal geometries of torsion angles in peptides, achieving competitive performance in the discussed tasks. In addition, to our best knowledge, we are the first to establish flow-matching models for torsional angles lying on torus manifolds and to propose a deep-learning model for target-specific peptide generation. (iii) **Dataset Establishment**. For training the deep learning model, we collect a benchmark dataset called `PPBench2024` consisting of 15593 protein-peptide pairs with high quality. Compared with previous protein-peptide datasets (Agrawal et al., 2019; Martins et al., 2023; Wen et al., 2018), `PPBench2024` augments and expands available data, and filters large amounts of low-quality data according to strict criteria, thus allowing deep-learning-based models to be trained on the datasets and to fulfill the peptide drug discovery tasks.

## 2. Background

### 2.1. Problem Statement

For a binding system composed of a protein-ligand pair (*i.e*. protein-peptide pair) as 𝒞, which contains *N*_pt_ amino acids of the protein and *N*_pp_ amino acids of the peptide, we represent the index set of the amino acids in the peptide as ℐ_pp_ = {1, …, *N*_pp_}, and the protein’s amino acids as ℐ_pt_ = {*N*_pp_ + 1, …, *N*_pp_ + *N*_pt_}. The amino acids can be represented by its type *s*^(*i*)^, atom coordinates 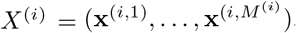, where *s*^(*i*)^ ∈ {1, …, 20}, **x**^(*i,j*)^ ∈ ℝ ^3^, and *M* ^(*i*)^ = *M* (*s*^(*i*)^), meaning that the number of atoms is decided by the amino acid types. The sets of 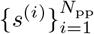 refers to a peptide sequence, and 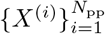 refers to its structure, which is the same for protein.

Therefore, 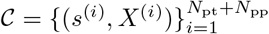 can be split into two sets as 𝒞= 𝒫 ∪ ℒ, where 𝒫 = {(*s*^(*i*)^, *X*^(*i*)^) : *i* ∈ ℐ_pt_} and ℒ = {(*s*^(*i*)^, *X*^(*i*)^) : *i* ∈ ℐ_pp_}. For *protein-specific peptide generation*, our goal is to establish a probabilistic model to learn the distribution of molecules conditioned on the target proteins, *i.e. p*(ℒ|𝒫).

### 2.2. Riemanian Flow Matching

Let 𝒫 = 𝒫 (ℳ) be the probability function space over a manifold ℳ with Riemanian metric *g. q* is probability distribution of data *x* ∈ ℳ, and *p* is the prior distribution. The probability path on ℳ as an interpolation in probability space written as *p*_*t*_ : [0, 1] →𝒫 satisfies *p*_0_ = *p* and *p*_1_ = *q. u*_*t*_(*x*) ∈ 𝒯_*x*_ℳ is the corresponding gradient vector of the path on *x* at time *t*. A Flow Matching (FM) tangent vector field *v*_*t*_ : [0, 1] × ℳ → ℳ is used to approximate *u*_*t*_(*x*) with the objective 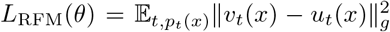, in which *θ* are the parameters in *v*_*t*_. *u*_*t*_ is intractable, and an alternative is to construct conditional density path *p*_*t*_(*x*|*x*_1_) whose gradient reads *u*_*t*_(*x*|*x*_1_), and use Conditional Flow Matching (CFM) objective as 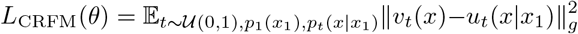, since FM and CFM objectives have the same gradients as shown in (Lipman et al., 2022; Chen & Lipman, 2023). Once the gradient field *v* is learned, one can use ordinary differential equations 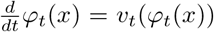 with *φ*_*0*_(*x*) = *x* ∼ *p*_0_ and *φ*_*t*_(*x*) = *x*_*t*_ to push the random variable *x*_*t*_ ∈ ℳ from prior distribution *p*_0_ to the data distribution *p*_1_.

## 3. Proposed Method

### 3.1. Peptide Structure Parameterization

There are several parameterizations for peptide structures. Here, we firstly focus on the four backbone atoms, *i.e*. {N, C*α*, C, O}. One effective parameterization follows AlphaFold2 (Jumper et al., 2021), in which the four atom positions *X*^(*i*)∗^ = **{x**^(*i*,1)^, …, **x**^(*i*,4)^} form a rigid body, and represent *X*^(*i*)∗^ ≜ (**x**^(*i*,2)^, *O*^(*i*)^) as variables on SE(3), in which **x**^(*i*,2)^ ∈ ℝ^3^ as C*α*’s coordinate is the translation vector and *O*_*i*_ ∈ SO(3) is an rotation matrix obtained by the ideal frame constructed by *X*^(*i*)∗^, leading 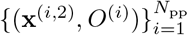 to represent the backbone structure. Hence, the structure representation lies on a manifold with 6*N*_pp_-degree of freedom. However, while the intra-residues’ internal geometries are fixed, *e.g*. bond lengths of ‘N-C*α*’ and ‘C*α*-C’, and bond angles of ‘N-C*α*-C’, the other inflexible inter-residues’ geometries of bond lengths such as ‘C-N’ and angles like ‘C-N-C*α*’ are not constrained in the parameterization.

In comparison, we focus on the torsion angles of *ϕ, ψ*, and *ω* (shown in Figure. 1) as redundant geometries according to the physicochemical conclusions (Padmanabhan, 2014) and our observations (See Appendix. B.2). While the local structures can be reconstructed with torsion angles and ideal bond lengths and angles with NeRF (Parsons et al., 2005), the global translation and orientation have to be determined to obtain the peptides’ poses relative to the target proteins (Corso et al., 2023). By this means, the backbone peptide structure can be represented as 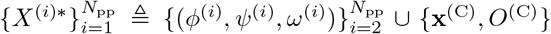. Note that *i* start from 2 because four consecutive atoms form a torsion angle, **x**^(C)^ is the global translation of the peptide centroid, and *O*^(C)^ is the rotation matrix of the global frame constructed by 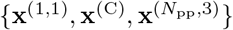. Hence, the backbone structure is parameterized by variables on 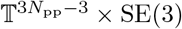. The advantages of the parametrization include (*i*) the inflexible bond lengths and angles are set to be constants, avoiding broken or overlapped bonds as unrealistic structures and (*ii*) fewer degrees of freedom to reduce the task difficulties (3*N*_bb_ + 3 < 6*N*_bb_).

**Figure 1:**
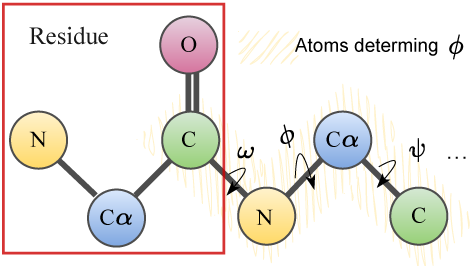
Backbone atom parameterization.

In the following, the CFM on the discussed manifolds will be described in Sec. 3.2 and Sec. 3.3, and for the sequence type CFM, it will be discussed in Sec. 3.4. Besides, for the other atoms in side-chains, we use the rotamer *χ*-angles as the parameterization, discussed in Sec. 3.8.

### 3.2. PPFlow on Torus

For each torsion angle lies in [− *π, π*), *N* torsion angles of a structure define a hypertorus 𝕋^*N*^, with *N* = 3*N*_pp_ − 3. To construct a conditional flow on 𝕋^*N*^, we follow RCFM, to obtain ***τ***_*t*_ using the geodesic connecting ***τ***_0_ and ***τ***_1_ with

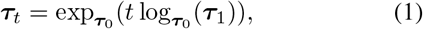

where ***τ***_0_ ∼ *p*_0_, ***τ***_1_ ∼ *p*_1_ and ***τ***_*t*_ ∈ 𝕋^*N*^, and the conditional vector field 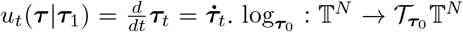 is the logarithm map, projecting points on 𝕋^*N*^ to the tangent space of 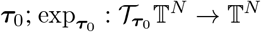 is the exponential map, projecting points back to torus (Chen & Lipman, 2023).

For the torus, we parameterize the manifold as the quotient space ℝ^*N*^ */*2*π*Z^*N*^, leading to the equivalence relations ***τ*** = (*τ* ^(1)^, …, *τ* ^(*N*)^) ≅ (*τ* ^(1)^ + 2*π*, …, *τ* ^(*N*)^) ≅ (*τ* ^(1)^, …, *τ* ^(*N*)^ + 2*π*) (Jing et al., 2023). By this means the prior distribution *p*_0_ is chosen as a product of standard to wrapped normal distribution on ℝ^*N*^ as

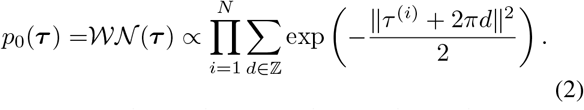

To construct the conditional path, considering the flow

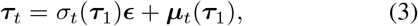

where *ϵ* ∼ 𝒲 𝒩(*ϵ*). To build the gradient vector, we propose that **Theorem 3**. in (Lipman et al., 2022) still holds for the defined path, as the following proposition:

#### Proposition 3.1

*Let p*_*t*_(***τ***|***τ***_1_) *be probability path as in Equation. 3. Its vector field has the form*

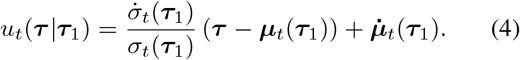

*Hence, u*_*t*_(***τ*** |***τ***_1_) *generates the probability path p*_*t*_(***τ*** |***τ***_1_).

The proof is given in Appendix. A.1. Besides, the following proposition ensures the correctness of the learning objective in Conditional Torus Flow Matching with proof in Appendix. A.2:

#### Proposition 3.2

*Given p*_*t*_(***τ***), ***τ*** ∈𝕋^*N*^, *the conditional and unconditional flow matching losses have equal gradients* w.r.t. *θ:* ∇_*θ*_*L*_UTFM_(*θ*) = ∇_*θ*_*L*_CTFM_(*θ*).

Following (Tong et al., 2023), we in practice construct our flow-matching objective in an Independent Coupling way:

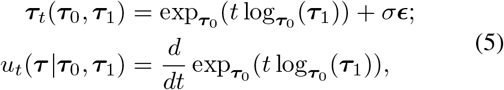

which is a specific instance of the above proposition. The explicit equations for geodesics on a hypertorus T^*N*^ can be complex and depend on the specific parametrization chosen for the hypertorus (Jantzen, 2012). Instead, we regard the torsion angles as mutually orthogonal and their interpolation paths on the torus are linear *w.r.t. t* in each direction. Therefore, we employ a fast implementation of calculating 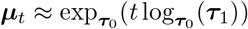 by

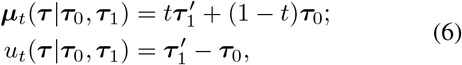

where ***τ*** ≅ ***τ***_1_^′^ = (***τ***_1_ − ***τ***_0_ + *π*) mod (2*π*) − *π*. The geodesic distance is approximated with the euclidean ones, as 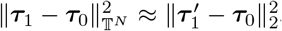, which satisfies definitions of premetric in Sec. 3.2 in (Chen & Lipman, 2023). This leads the closed-from expression of the loss to train the conditional Torus Flow Matching to

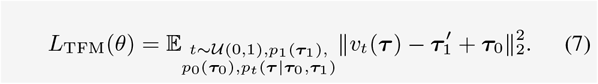

### 3.3. PPFlow on SE(3)

The torsion angles can reconstruct the local structures of the peptides, while in the global coordinate system, to represent the positions of residues, we need to determine their global translations and rotations.

As discussed in Sec. 3.1, the pose representation on SE(3) can be decomposed into global translation as **x**^(C)^ ∈ ℝ^3^ and rotation *O*^(C)^ ∈ SO(3). For notation simplicity, we omit the superscript ^(C)^ in this part. To model the probability path of **x**_*t*_, we employ vanilla Gaussian CFM on Euclidean manifolds, with Independent Coupling techniques:

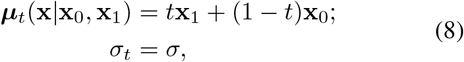

where **x**_0_ ∼ 𝒩(**x** |**0**, *I*), thus leading to the Gaussian probability path of *p*_*t*_(**x x**_0_, **x**_1_) = 𝒩(**x** |***μ***_*t*_, *σ*_*t*_), and the loss to train the conditional Euclidean Flow Matching as

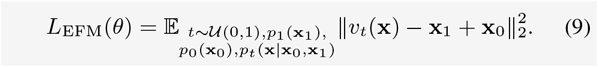

To model the probability path of rotation matrix *O*_*t*_ ∈ SO(3), we employ SO(3)-CFM (Yim et al., 2023a), in which 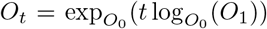. In the implementation, since the SO(3) is a simple manifold with closed-form geodesics, the exponential map can be computed using Rodrigues’ formula and the logarithmic map is similarly easy to compute with its Lie algebra so(3) (Yim et al., 2023b). The prior distribution to sample *O*_0_ is defined as isotropic Gaussian distribution, by first parameterizing *O*_0_ in axis-angle, where the axis of rotation is sampled uniformly and the density function of rotation angle *ϑ* reads 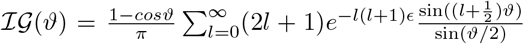 (Creasey & Lang, 2017). For the conditional gradient field, we employ fast numerical tricks (Bose et al., 2023) to calculate the SO(3) component of the global Orientation conditional Flow Matching objectives:

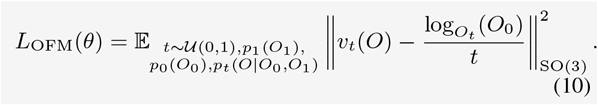

The correctness of the learning objective of Equation. 9 and 10 is proposed and proven in (Lipman et al., 2022) and (Bose et al., 2023), respectively.

### 3.4. PPFlow on Amino Acid Types

For the amino acid sequence 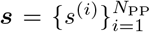, we directly model the probability vector of each type, where 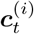 is defined as the probability vector of multinomial distribution with 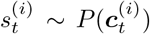. We omit ^(*i*)^ for notation simplicity. To build a path, we define ***c***_1_ = onehot(*s*_*i*_), and 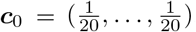. Analogously to the Euclidean flow, we define ***c***_*t*_ = *t****c***_1_ + (1 −*t*)***c***_0_, and *u*_*t*_(***c***|***c***_0_, ***c***_1_) = ***c***_1_ −***c***_0_. It is easy to prove ***c***_*t*_ is a probability vector since its summation equals 1 across all types. Instead of an MSE training loss in R^20^, we, inspired by multinomial diffusion (Hoogeboom et al., 2021), propose a multinomial flow matching objective:

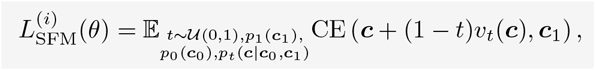

where CE(·,·) is cross-entropy, and the loss directly measures the difference between the true probability and the inferred one ***ĉ***_1_ = ***c*** +(1 − *t*)*v*_*t*_(***c***). For the whole sequence, the loss function is a summation of 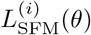 over *i*, as

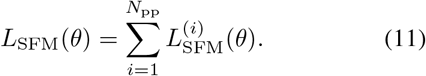

### 3.5. Overall Training Loss

The overall loss function is the summation of the four loss functions in Equation. 7, 9, 10 and 11. To fully utilize the sufficient contextual information as conditional inputs, for each flow-matching vector field *v*_*t*_(·), we instead use all the structure-sequence contexts as input. In this way, we will first sample 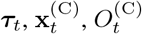 and 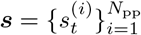 through the defined conditional probability paths, and then employ NeRF (Parsons et al., 2005) to reconstruct the sequence and backbone structure in local frame, and translate and rotate the structures such that its centroid coordinate and orientation equals to 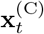 and 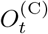. Thus, we can obtain 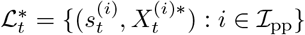. Finally, the whole binding complex at *t* is used as inputs of *v*_*t*_(·), as 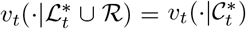. Therefore, our overall training loss is written as

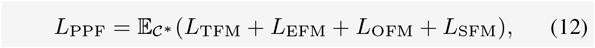

in which 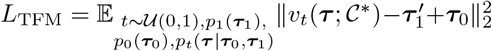, and the other three can be written in a similar way. We write 𝒞^∗^ = rec(***τ***, **x**^(C)^, *O*^(C)^, ***s***), meaning the input binding complexes are reconstructed from the four variables.

### 3.6. Sampling with ODE

For the trained flow-matching vector fields, we sampling process is the solution of the following ordinary the differential equation as 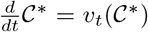, in which the defined method is employed as the numerical solution:

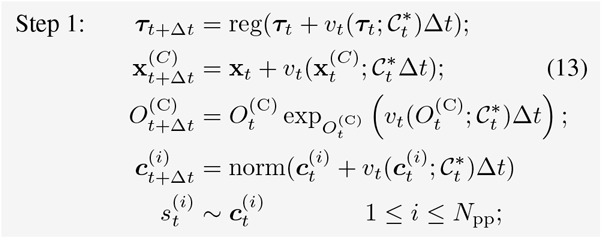

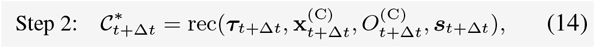

where norm(·) means normalizing the vector to a probability vector such that the summation is 1, and reg(·) means regularize the angles by reg(***τ***) = (***τ*** + *π*) mod (2*π*) − *π*.

### 3.7. Parameterization with Neural Networks

#### Encoder

We first adopt two multi-layer perceptrons (MLPs). The embedding of single residues is obtained by one MLP, which encodes the residue type, backbone dihedral angles, and local atom coordinates. The other MLP for residue pairs encodes the distance, the dihedral angles between the *i* and *j* residues, and their relative positions. Then, we stack 6-layer transformers with self-attention mechanisms used to update the inter-residues’ embedding and obtain residues’ embedding for each amino acid by ***h***_*i*_ ∈ ℝ^*D*^.

#### Equivariance

The conditional distribution of *p*(ℒ) have to be roto-translational equivariant to ensure the generalization, *i.e. p*(*O* ℒ + **x** | *O* ℛ + **x**) = *p*(ℒ |ℛ), for **x** ∈ R^3^ and *O* ∈ SO(3), where *O* + **x** = (*s*^(*i*)^, *OX*^(*i*)^ + **x**). To avoid the problem of non-existence of such a probability due to translation, we here adopt the zero-mass-center techniques (Rudolph et al., 2020; Yim et al., 2023b), by subtracting the mass center of the conditional receptor from all inputs’ coordinates to the neural network (Guan et al., 2023; Lin et al., 2022; Satorras et al., 2021), which also helps to improve the training stability. Further, in terms of CFM, the following proposition indicates the roto-translational equivariance of each flow-matching vector field.

##### Proposition 3.3

*Let p*_0_(***τ***), *p*_0_(**x**^(C)^), *p*_0_(*O*^(C)^), *and p*_*t*_(***c***) *be SE*(*3*)*-invariant distribution, and the flow-matching vector field 𝒗*_*t*_(**x**^(C)^|𝒞^∗^) *be SE*(*3*)*-equivariant, 𝒗*_*t*_(*O*^(C)^ |𝒞 ^∗^) *be T*(*3*)*-invariant and SO*(*3*)*-equivariant, and 𝒗*_*t*_(***c*** |𝒞^∗^) *and 𝒗*_*t*_(***τ*** |𝒞^∗^) *be SE*(*3*)*-invariant, then the density p*(ℒ ^∗^ |ℛ) *generated by the ODE sampling process SE*(*3*)*-equivariant*.

We employ ITA (Jumper et al., 2021) with LoCS (Kofinas et al., 2022) (For details see Appendix. A.4), to ensure the equivariance and the invariance of the vector fields in 3.3.

### 3.8. Side-Chain Packing

For the complete solution to full-atom design, after the backbone atoms are generated, we employ a Rotamer Density Estimator (RDE) (Luo et al., 2023) as the probabilistic model to model the side-chain rotamers 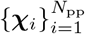. It uses a Conditional Flow on 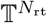 based on the rational quadratic spline flow (Durkan et al., 2019; Rezende et al., 2020), where *N*_rt_ is the total rotamer number. In practice, we use the pre-trained version of RDE capable of perceiving the side-chain conformations by training the model on large datasets of `PDB-REDO` (Joosten et al., 2014). Further, we conduct fine-tuning on our protein-peptide complex datasets. The effectiveness of the side-chain packing model named RDE-PP, transferred from protein-protein complexes to the protein-peptide ones, is empirically shown in Sec. 5.6.

The overall workflow of PPFlow including the backbone generation and side-chain packing is shown in Figure. 2.

**Figure 2:**
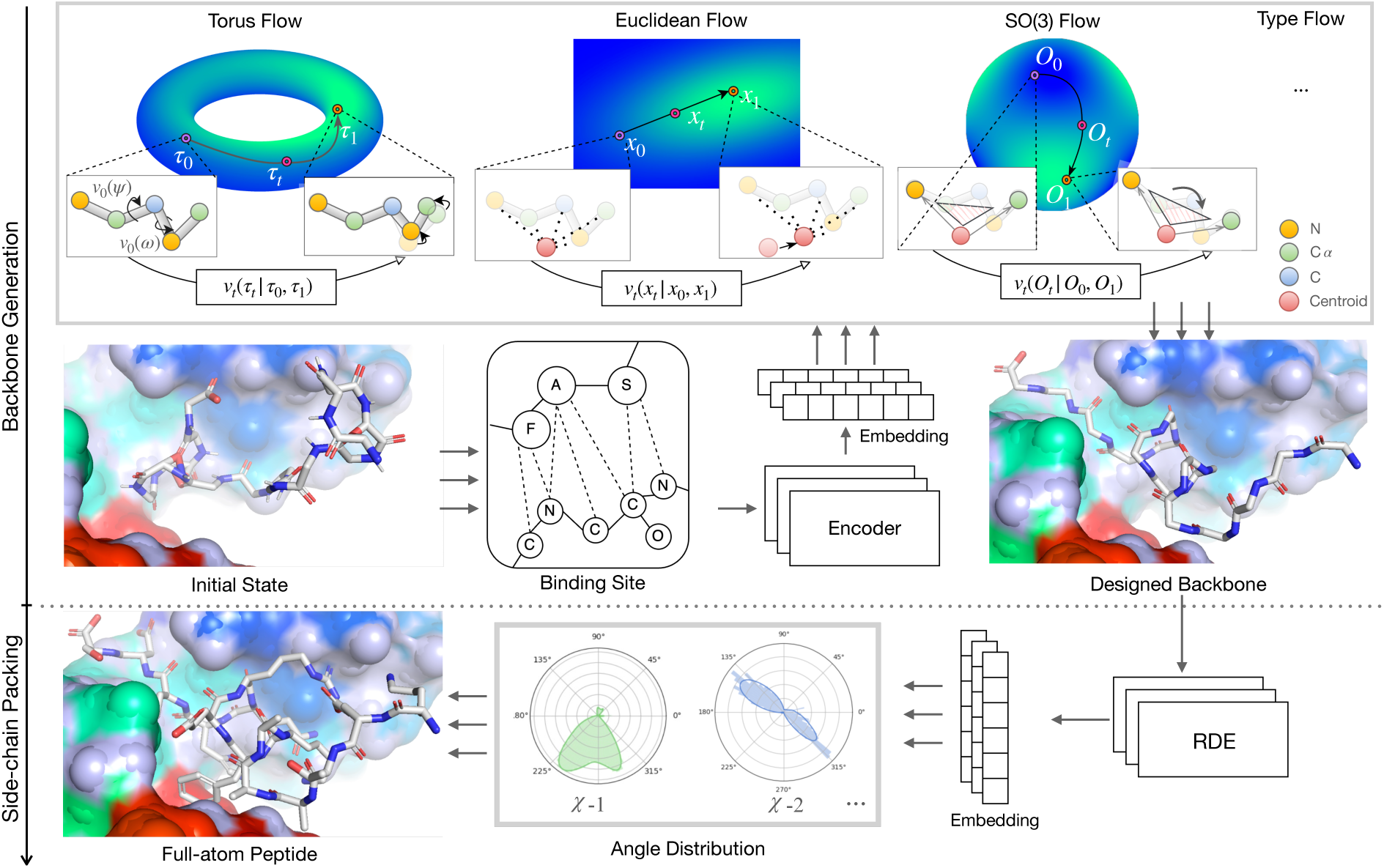
Workflows of PPFlow in target-aware peptide drug generation task.

## 4. Related Work

### Structure-based drug design

Success in 3D molecule generation and increasing available structural data raises scientific interest in structure-based drug design (SBDD). Grid-based methods regard the Euclidean space as discrete and predict the molecules’ structures on the grids (Luo et al., 2021; Masuda et al., 2020). With great advances in Graph Neural Networks (Wu et al., 2021; Liu et al., 2021; Wu et al., 2022c), Equivariant neural networks have advanced the structure predictions and helped to improve the tasks greatly (Peng et al., 2022b; Liu et al., 2022). Then, diffusion methods (Guan et al., 2023; Schneuing et al., 2022; Lin et al., 2022) attempt to generate the ordered atoms’ positions and types at full atom levels. Further, a series of works of fragment-based drug design are proposed, by mimicking classical drug design procedures to generate both atoms and functional groups that form molecule drugs (Zhang et al., 2023b; Lin et al., 2023).

### Protein generation

Machine learning methods for protein modeling have achieved great success in recent years (Huang et al., 2023; Wu et al., 2024; 2022b; Wu & Li, 2023), and peptides can be regarded as fragments making up a protein. Techniques on protein backbone generation assist realistic peptide structure design. For example, FoldingDiff (Wu et al., 2022a) is also based on redundant angle generation, which employs diffusion models on torus to achieve backbone design. RFDiffusion and Chroma (Watson et al., 2023; Ingraham et al., 2022) use different diffusion schemes, and achieve state-of-the-art generation performance on protein generations. Recently, protein backbone generation methods employ flow matching techniques, to explore the applicability and effectiveness (Yim et al., 2023a; Bose et al., 2023). For protein side chains, methods usually focus on protein-protein complexes, such as RED-PPI (Luo et al., 2023) and DiffPack (Zhang et al., 2023a). Our side-chain packing methods follow RDE-PPI, and achieves a good generalization performance on peptides.

## 5. Experiment

### 5.1. Dataset

#### Training set

To satisfy the need for massive data to train deep learning models, we construct `PPBench2024`, through a series of systematic steps: *First*, we source complexes from the RCSB database (Zardecki et al., 2016), specifically selecting those containing more than two chains and excluding any with nucleic acid structures, and defining interactions between a pair as a minimum intermolecular distance of 5.0Å or less. *Subsequently*, only those complexes featuring peptide chains that do not exceed 30 amino acids in length are included, to better mimic existing peptide drug sizes. *Then*, the water molecules and heteroatoms are eliminated. *Finally*, we select only peptide molecules composed entirely of amino acids, remove the modified peptides with functional groups other than amino acids, and filter peptides with broken bonds according to the ideal bond length, *i.e*. the bond is unbroken if the observed bond length is between the ideal length plus or minus 0.5Å. The number of final screened complex instances of peptide-protein pairs is 9070. Further, we screen the existing datasets of `PropediaV2.3` (Martins et al., 2023) and `PepBDB` (Wen et al., 2018) with the same criterion, leading to additional 6523 instances to expand it. Appendix. B.1 gives details. We split the `PPBench2024` into training and validation sets according to the clustering of the proteins that are closest to the peptide ligand via MMSeqs2 (Steinegger & Söding, 2017) with the ratio of 9 : 1.

#### Test set

To evaluate the model performance, we use an existing benchmark dataset called `PPDBench` (Agrawal et al., 2019) consisting of 133 pairs with peptides’ lengths ranging from 9 to 15 as the test set. We eliminate all complexes from `PPBench2024` whose PDB-ID are the same as those in `PPDBench` to avoid potential data leakage.

### 5.2. Baseline Models

Our method has been opened to the public in https://github.com/Edapinenut/ppflow. For comparison, we extend two models for the following tasks, including

i. DiffPP, as a variant of DiffAB, which is a diffusion model for generating CDRs in antibodies targeted at antigens. DiffPP parametrizes the protein backbones in the same way as AlphaFold2, in which the positions of atoms in a residue are determined by C*α*’s translation vector **x**_*i*,2_ and the rotation matrix *O*_*i*_ ∈ SO(3) of the frame constructed by positions of {N, C*α*, C}. The diffusion and reverse process on translation and orientation variables are modeled by DDPM (Ho et al., 2020) and SO(3)-DPM (Leach et al., 2022). For amino acid types, it uses a multinomial diffusion (Hoogeboom et al., 2021).
ii. DiffBP-PP, as a variant of DiffBP or TargetDiff, as an atom-level diffusion model for generating molecules that bind to specific proteins. It decomposes the residues into atoms, and uses DDPM to model the distribution of atoms’ positions and D3PM (Austin et al., 2021) to predict amino acid types through mask language modeling techniques.

These two models are the state-of-the-art models for target-specific protein and molecule generation, respectively. For other classical models, we describe them in different tasks.

### 5.3. Peptide Generation

#### Metrics

We choose 6 metrics for evaluating the quality of generated peptides. **∆*G*** is binding energies calculated by ADCP re-docking (Zhang & Sanner, 2019), since ADCP has shown the best performance in estimating the binding energies and poses of protein-peptide complexes (Weng et al., 2020). ∆*G* reflects the potential of a peptide being pharmaceutically active towards the targets, and **IMP%-B** gives the percentages of the designed peptides with lower (better) ∆*G* than the reference peptides. The mean of reference ∆*G* is −427.72kcal*/*mol. Stability evaluates binding scores of the original pose of the generated peptides, which directly reflects the quality of generated peptides without re-docking. The stability scores are calculated by FoldX (Schymkowitz et al., 2005) since it performs fast and accurate stability calculation in protein binding tasks compared with other energy-based methods (Luo et al., 2023). We give **IMP%-S** as the percentages of the designed ones with better stability than reference rather than average because some extremely unstable structures will make the comparison on average values meaningless. **Validity** is the ratio of the designed peptides that is chemically valid, through the criterion of whether the bonds of the atoms that should be bonded are broken. We follow our filtering process of datasets and define the bond is not broken if its length is within 0.5Å above and below the ideal value. **Novelty** is measured considering two aspects of structure and sequence (i) the fraction of peptides with TM-score < 0.5 as used in (Lin & AlQuraishi, 2023) and (ii) the sequence overlap (SeqOL) less than 0.5. The given score is the mean ratio of novel peptides *v.s*. references. **Diversity** is the product of pairwise (1 −TM-score) and (1 −SeqOL) of the generated samples averaged across their target proteins. The higher the score, the more diverse peptides a model generates. Note that all the metrics except Validity are measured with chemically valid peptides.

#### Setup

For each model for comparison, we generate 20 peptides for each conditioned protein because the evaluation process requires excessive time. For example, ∆*G* obtained by ADCP requires more than 10 minutes for one pair of protein and peptides on a server with 128 CPU threads. Besides, since the greater the number of atoms, the more likely it is that binding interactions will occur, we eliminate the effects of peptide sizes (lengths) on the binding affinity by setting the generated sequence lengths the same as references.

#### Results

Table. 1 gives the comparison of peptide generation task with generated examples shown in Figure. 4. For PPFlow, we test two variants of it. PPFlow-BB (PPFlowBackBone) uses the generated backbone structures with Rosetta side-chain packing (Alford et al., 2017) before re-docking scoring and stability calculation, and side-chain atoms in PPFlow-FA (PPFlow-FullAtom) is predicted by RDE. Note that the backbone structures before side-chain packing are the same for the two variants, leading to metrics except IMP%-S being of little difference. Besides, for the others, the side-chain atoms are constructed by Rosetta. It can be concluded that (i) PPFlow has the greatest potential to generate peptide drugs with high binding affinity towards the target protein according to ∆*G* and IMP%-B metrics. (ii) The complexes of peptides of original poses generated by PPFlow and the target proteins are the most stable, with the highest IMP%-S metrics. (iii) PPFlow and DiffPP generate a comparably high ratio of novel peptides, while peptides generated by PPFlow are more diverse. (iv) The rotamers estimated by RDE-PP are more stable in the binding site than Rosetta according to IMP%-S metrics of PPFlow-BB and PPFlow-FA. (v) With the bond lengths and angles fixed, the structures generated by PPFlow are chemically valid, while DiffBP-PP shows the lowest Validity due to the highest degree of freedom it models, *i.e*. 4 × 3*N*_pp_, for atom-level backbone structure, which also empirically shows the infeasibility of SBDD methods to generalize to peptide drug design. Therefore, in the following parts, we will not compare DiffBP-PP as baselines.

**Table 1:** Comparison for target-aware peptide generation. Values in **bold** are the best.

| | $\Delta G(\downarrow)$ | IMP%-B(↑) | IMP%-S(↑) | Validity(↑) | Novelty(↑) | Diversity |
| --- | --- | --- | --- | --- | --- | --- |
| DIFFPP | -319.54 | 28.75% | 7.56% | 0.94 | 0.79 | 0.28 |
| DIFFBP-PP | -221.06 | 19.34% | 4.09% | 0.42 | 0.76 | 0.41 |
| PPFLOW-BB | -349.59 | **36.02%** | 10.34% | **1.00** | **0.84** | **0.76** |
| PPFLOW-FA | **-351.27** | 35.43% | **12.88%** | **1.00** | **0.84** | **0.76** |

### 5.4. Peptide Optimization

#### Setup

We apply our model to another common pharmaceutical application: optimization of the existing peptides. To optimize a peptide, we first choose a start time start time ∈ [0, 1], and use the probability paths constructed in Sec. 3.2, 3.3 and 3.4, to sample a perturbed peptide 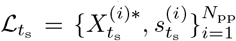. By using the ODE sampling methods in Sec. 3.6, the trained PPFlow iteratively updates the sequences and structures to recover the peptides as a set of optimized ones. For DiffPP, we use the same setting as (Luo et al., 2022). The ‘num_step’ is the number of how many optimization steps used in the generative diffusion process, where the start step of noised peptides is (total_step− time_step). The conversion between them is total_step −num_step = total_step × start time. We sample 20 new peptides for each protein receptor in `PPDBench` and evaluate them in the same metrics as discussed in Sec. 5.3. We set total step = 100 in this part.

#### Results

Figure. 3 gives the comparison on peptide optimization results. It can be concluded that (i) The larger number of optimization steps contributes little to the improvement in the binding affinity, but PPFlow usually generates more peptides with higher binding affinities; (ii) Optimized peptides are usually similar to the original one in small optimization steps since the novelty are smaller, which is desired in many practical applications; (iii) The diversity is more controllable in PPFlow by changing the number of optimization steps.

**Figure 3:**
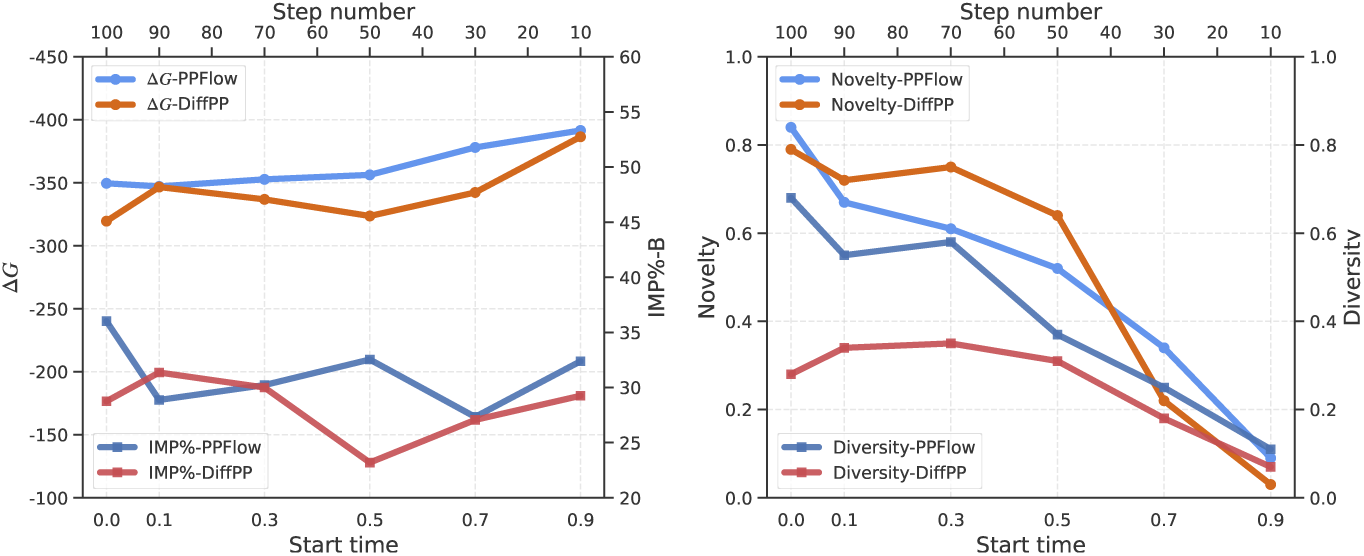
Metrics for generated peptides of methods in different optimization steps.

**Figure 4:**
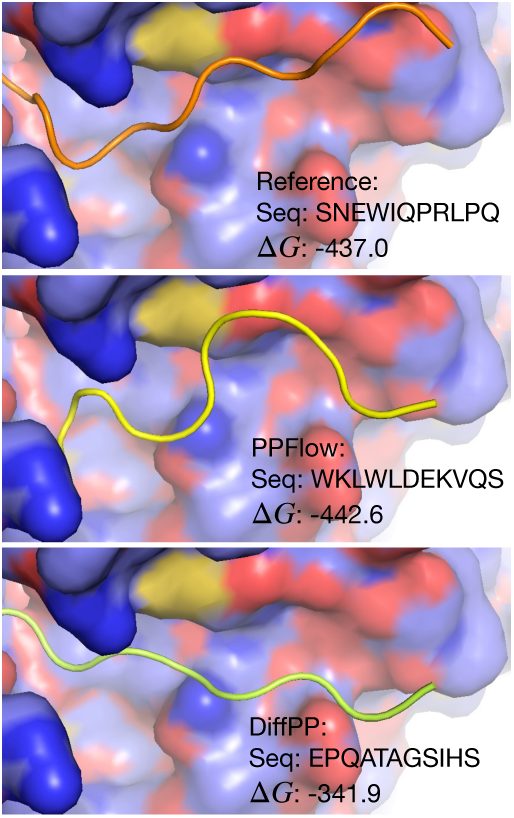
Peptides designed by different methods and reference.

### 5.5. Protein-Peptide Docking

Besides the two tasks, in this part, we generalize our model to the peptide flexible re-docking task to figure out if the model can learn the binding poses of the ligands. The task can be regarded as establishing a probabilistic model of 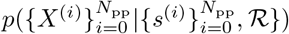. For DiffPP and PPFlow, 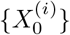, as the initialized peptide structures, are obtained by first moving them to the pocket center, and then adding Gaussian noise to each atom. We retrain DiffPP and PPFlow with the sequences given as additional conditions. For classical docking methods, we choose HDock (Yan et al., 2020) which is a re-docking method for protein-protein interactions, and VinaDock (Eberhardt et al., 2021) proposed for molecule docking. We choose the best 10%/30%/50% poses for comparison on ligand-RMSD (**L-RMSD**) between C*α* atoms and centroid-RMSD (**C-RMSD**), and the success rate (**Success%**) of the docked ligands’ RMSD smaller than 2/4/8Å. It can be concluded in Table. 2 that (i) DiffPP shows the best docking performance on L-RMSD and Success% because it directly optimized the RMSD between C*α* atoms, and PPFlow models the ligands’ centroid most accurately. (ii) The deep-learning-based generative models can achieve competitive performance with the classical energy-based ones, while the latter requires enormous rounds of iteration, costing dozens of times longer for computation.

**Table 2:** Comparison for flexible peptide re-docking.

| Methods | L-RMSD(↓) |  |  | C-RMSD(↓) |  |  | Success%(↑) |  |  | Time (s) |
| --- | --- | --- | --- | --- | --- | --- | --- | --- | --- | --- |
|  | 10% | 30% | 50% | 10% | 30% | 50% | 2 | 4 | 8 |  |
| DIFFPP | **8.01** | **15.26** | 28.47 | 6.35 | 12.68 | 30.25 | **9.24** | **20.21** | **34.45** | 27 |
| PPFLOW | 12.44 | 17.82 | 31.24 | **2.60** | **3.28** | **4.26** | 6.13 | 11.24 | 22.47 | 46 |
| HDOCK | 9.95 | 18.65 | **23.07** | 7.29 | 10.65 | 18.84 | 6.40 | 10.72 | 25.50 | 186 |
| VINA | 11.38 | 19.03 | 27.69 | 6.27 | 9.56 | 16.59 | 5.45 | 9.96 | 21.12 | 1325 |

### 5.6. Side-Chain Packing

To figure out the effectiveness of our fine-tuned model of RDE-PP, We compare it with two baseline methods Rosetta(fixbb) (Leman et al., 2019) and the unpretrained version RDE-PP(w/o pt). For each reference peptide, we initialize the rotamers randomly with uniform angle distribution, and sample 20 conformations on side chains. We evaluate the mean absolute error (**MAE**) of the predicted sidechain torsional angles as MAE = 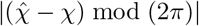 where 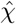 is the estimated ones and *χ* is the ground truth, and negative likelihood (**NLL**). In Table. 3, the results demonstrate that the RDE-PP with pretraining outperforms the baselines on three of the four torsional angles in terms of the evaluated metrics.

**Table 3:**
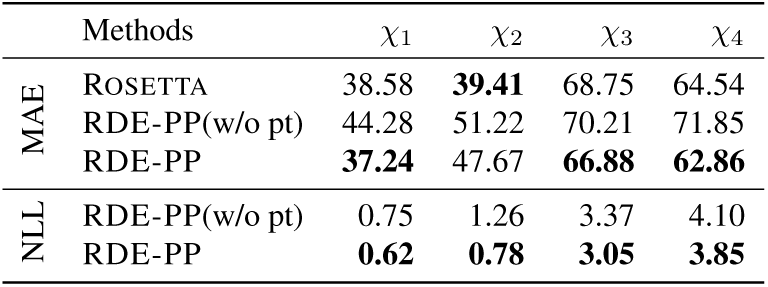
Comparison for side-chain packing.

## 6. Conclusion

In this paper, we focus on the target-specific peptide design tasks. To fulfill it, we first establish a dataset called `PPBench2024`, and then propose a flow-matching generative model on torus manifolds called PPFLOW, which attempts to learn the distribution of torsion angles of peptide backbones directly. Experiments are conducted to evaluate its performance on several tasks, indicating its superior performance on target-aware peptide design and optimization.

However, since the classical docking methods for binding pose calculation are extremely slow, we extend our model to docking tasks. Still, it is not as competitive as the baseline models, which will be our future focus. Besides, the performance gaps between the extended side-chain packing model and the classical ones are still small, urging us to develop a side-chain packing model with high prediction accuracy.

## Supporting information

appendix with proof, method details and experiments

ppflow workflow

## Acknowledgements

This work was supported by the Science & Technology Innovation 2030 Major Program Project No. 2021ZD0150100, National Natural Science Foundation of China Project No. U21A20427, Project No. WU2022A009 from the Center of Synthetic Biology and Integrated Bioengineering of Westlake University, and Project No. WU2023C019 from the Westlake University Industries of the Future Research. Finally, we thank the Westlake University HPC Center for providing computational resources. Besides, we thank the help of Dr. Tailin Wu, and his great efforts in his deep insights into cutting-edge issues, the guidance provided in the rebuttal, and the funding that supplied us with the necessary equipment for our research.

## Impact Statement

With the long-term continuation of COVID-19, more and more AI scientists are beginning to turn their research interests to drug design. The focus of this paper is on one of them – peptide drugs. There is very limited work aimed at using AI algorithms to design and optimize peptide drugs. However, peptide drugs are now rising as effective therapeutics, with hundreds of them proving to be successful, such as semaglutide for obesity and diabetes. Here, this work presents a considerable contribution to establishing a large dataset and deep-learning model for protein-specific peptide design, in which the effectiveness is validated empirically. As far as we know, it is the first AI-assisted peptide drug design solution, so we here emphasize the social significance of the work and hope to get the attention of the reviewers and the review committee.

## A. Method

### A.1. Proof of Proposition. 3.1

By Equation. 3, we can obtain 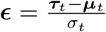. Besides, from the definition of 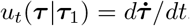, it can be written as

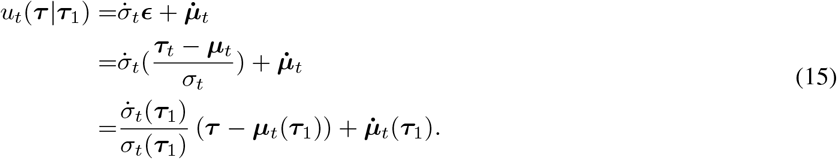

### A.2. Proof of Proposition. 3.2

Firstly, we claim the disintegration of measures of *p*_*t*_(***τ***), as *p*_*t*_(***τ***) =П_*i*_ *p*(*τ* (*i*)). For *p*_0_(***τ***), the disintegration satisfies. For *p*_1_(***τ***), our parametrization assumes the torsion angles are orthogonal and the distribution is independent, so it also satisfies disintegration. In this way, it is easy to obtain an intermediate probability *p*_*t*_ satisfies disintegration, and similar for the conditional probability *p*_*t*_(***τ*** |***τ***_1_).

Then we can factorizes the metric on 𝕋^*N*^ into S × · · · × S, and *p*(*τ*) ∈ P(S).

Let 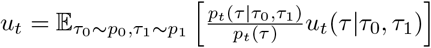, and

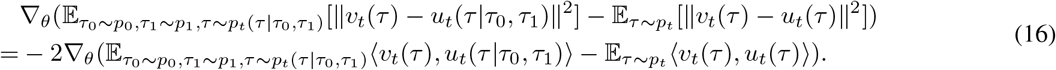

Then,

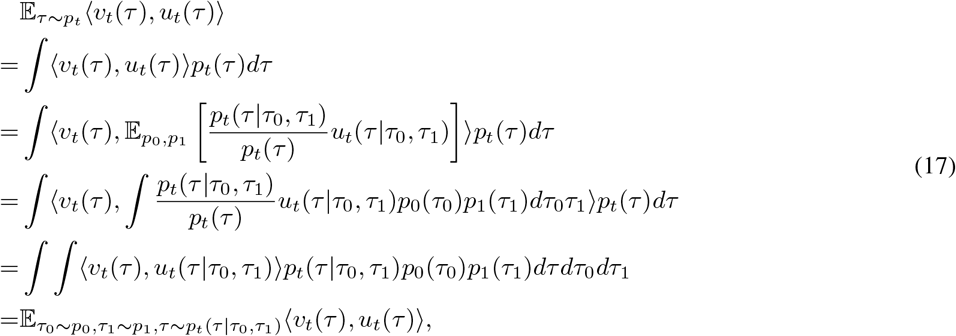

where in the last equality we change the order of integration. It proves Equation. 16 equals 0.

### A.3. Proof of Proposition. 3.3

The proof is inspired by (Köhler et al., 2020; Lin et al., 2023), as follows:

#### Lemma A1

Let T_*g*_(·) be the operation in SE(3), if the following update function in the ODE sampler (Sec. 3.6) for the atom level’s positions are defined as

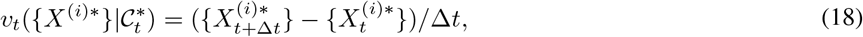

in which 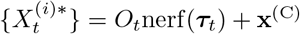. The invariance and equivariance of the following functions in the updating process are listed as and the prior distribution as

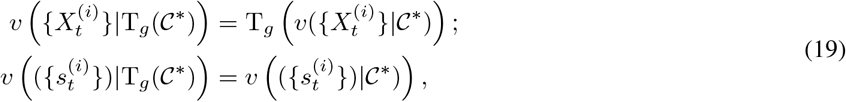

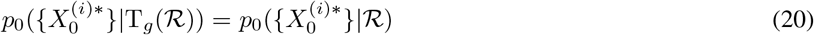

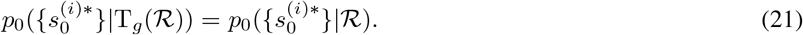

Then the distribution which the final structures and sequences are sampled from as *p* (L|R) is SE(3)-equivariant.

**Proof:** In the following, we write 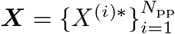, and 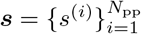 for notation simplicity. For the defined *𝒗*_*t*_, we first obtain that the update process is equivariant, as

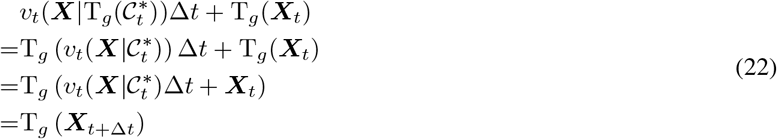

and for each transition kernel 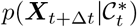, it is SE(3)-equivariant, since

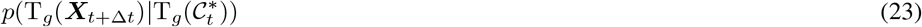

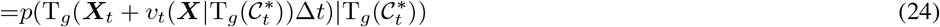

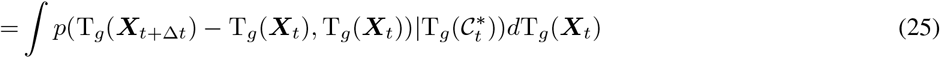

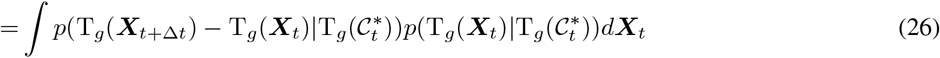

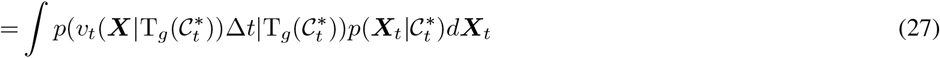

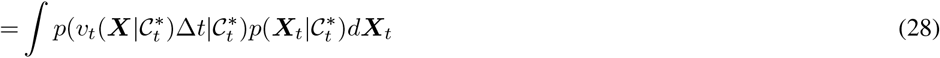

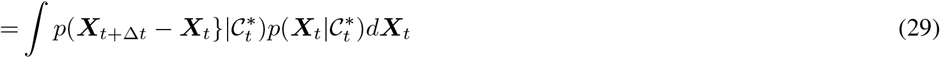

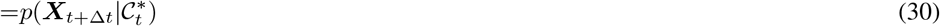

And it is easy to obtained that the transition kernel 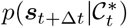 is SE(3)-invariant, since 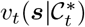 is invariant. Besides, for 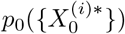 and 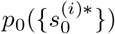, it is a SE(3)-invariant distribution. Therefore,

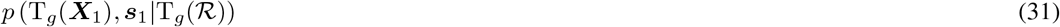

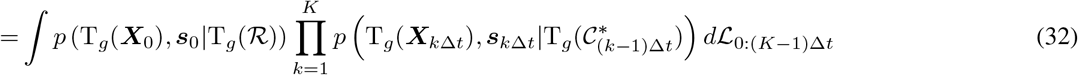

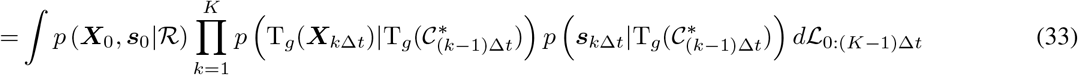

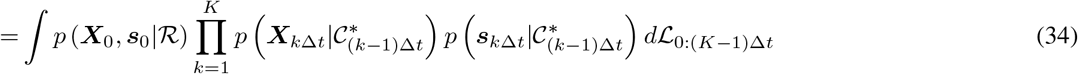

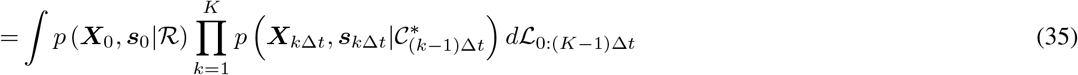

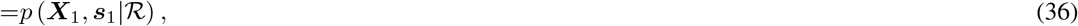

where *K* is the step number of the ODE sampler, and Δ*t* = 1*/K*. Because *p* (T_*g*_(***X***_1_), ***s***_1_|T_*g*_(R)) = *p* (***X***_1_, ***s***_1_|R), which is equivalent to 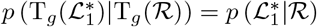, the Lemma is proved.

Then we write 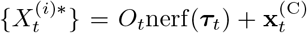, and the following proposition can give the conditions that the transition kernels of 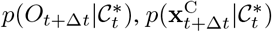 and 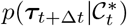 and 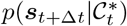 should satisfiy.

#### Lemma A2

If 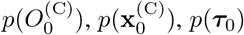 and *p*(***s***_0_) are SE(3)-invariant, and 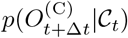 is SO(3)-equivariant and T(3)-invariant, 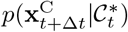 is SE(3)-equivariant, 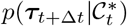 and 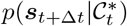 is SE(3)-invariant, then 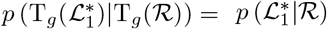, where 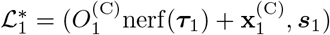.

**Proof:** Here for T_*g*_, we can decompose it as T_*g*_ = T_*r*_ ○ T_*t*_, meaning the rotation and translation opeartions as SE(3) ≅ SO(3) + T(3). By this mean,

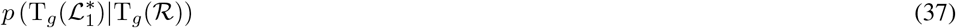

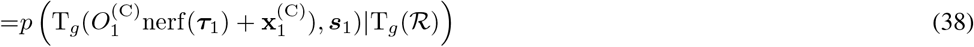

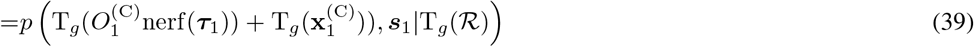

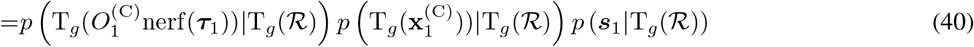

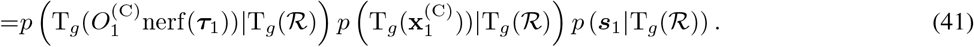

Because nerf(***τ***_1_) is always reconstructed with unit rotation diag(1, 1, 1) and zero-mass-centered, therefore,

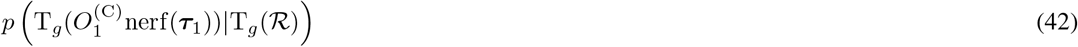

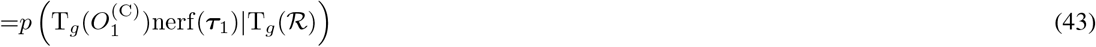

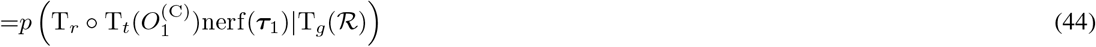

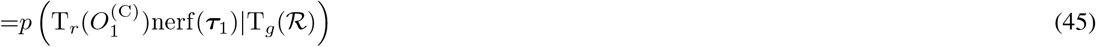

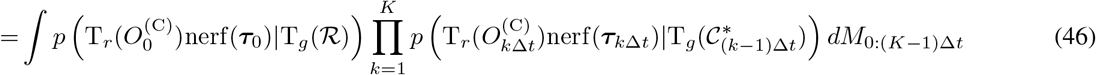

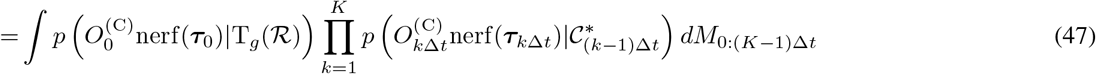

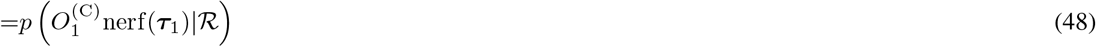

where 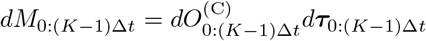. For 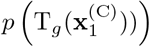 and *p* (***s***_1_|T_*g*_(ℛ)), the equivariance and invariance are also easy to obtain. Therefore, the Lemma A2. is proved.

Finally, **PROOF OF PROPOSITION 3.3**:

For the transition kernels which are generated by the following equations as

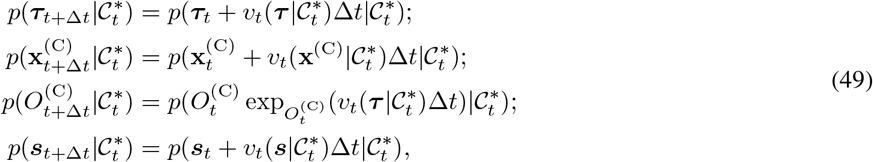

if the conditions of flow-matching vector fields in **Propostion 3.3** are all satisfied, then 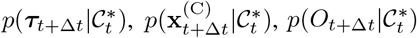 and 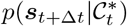 satisfies the conditions in **Lemma A.2** resepectively, and 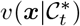 satisfies **Lemma A.1**. From both perspectives, the roto-translational equivariance can be proved.

We demonstarte the first proof path, to shown that the conditions of **Lemma A.2** holds.

For ***τ*** :

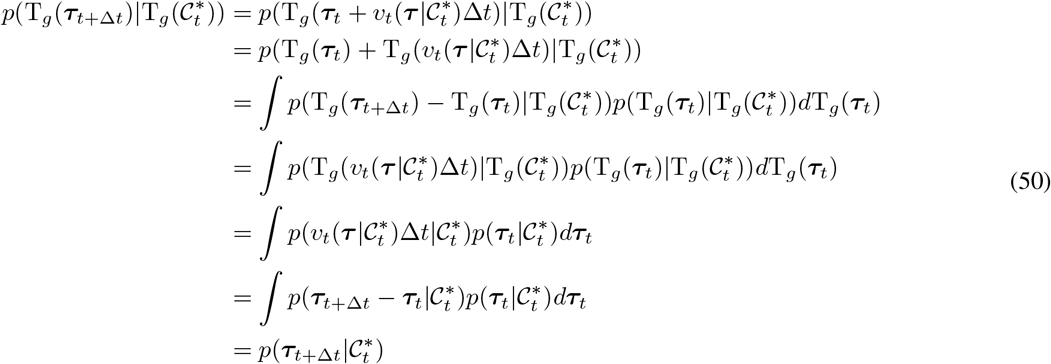

Beside, for the other three variables, the equations can be deduced similarly.

### A.4. LOCS Updating

Because the outputs should include the gradient vectors of rotation angle, global translational vector, rotation matrix, and type probability, we employ the LOCS to output the vector fields by

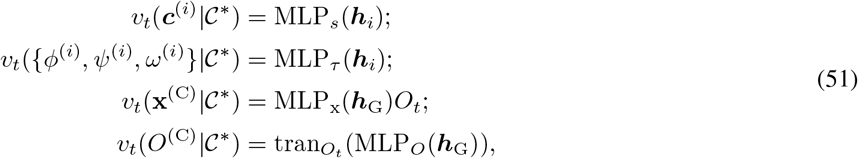

in which ***h***_G_ = ∑_*i*_ ***h***_*i*_ is the global representation obtained by summation, MLP_*s*_ : ℝ^*D*^ → ℝ^20^, MLP_*τ*_ : ℝ^*D*^ → 𝕋^3^, MLP_x_ : ℝ^*D*^ → ℝ^3^ and MLP_x_ : ℝ^*D*^ → ℝ^3^. MLP_*O*_ predicts a vector in Lie group so(3), and translate it to *O*_*t*_’s tangent space by 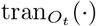. The output vector fields satisfy the equivariance and invariance conditions in Proposition. 3.3.

## B. Dataset Statistics

### B.1. Data preprocess

The construction of raw `PPBench2024` is given in the Sec. 5.1. For the additional complexes from `PropediaV2.3` and `PepBDB`, we give the detailed screening process in the Figure. 5. Because in PropediaV2.3, one peptide can be paired with several chains in a protein complex, another re-matching step should be conducted first.

Then, we give an empirical distribution on the peptide lengths of `PPBench2024`, in Figure. 6(a).

### B.2. Analysis on Geometry

As the PPFlow models the internal redundant geometry of the peptides, here we give a statistical illustration to show the flexible geometries that need to be generated. NeRF can use the following geometries in Figure. 6(b), 6(c) and 6(d) to reconstruct the full backbone structures. For these internal geometries, it can be concluded that the torsion angles of ‘N-C*α*-C-N’ and ‘C-N-C*α*-C’ are the two most flexible geometries. Besides, ‘C*α*-C-N-C*α*’ is theoretically inflexible since the constraints on peptide bond. However, in the observation, we find that it will deviate from the ideal value (*π*) a lot (about plus and minus 7°). In this way, we include it as another flexible geometries that the model needs to generate. These three torsion angles are named *ϕ, ψ* and *ω*, respectively in formal definition.

### B.3. Experiment

Here we give the hyper-parameters and other training details. The learning rate *lr* is 5*e* − 5. In all training, the max training iteration is 200000. `LambdaLR` schedule is used, with `lr lambda` is set as 0.95 × *lr*. The batch size is set 16 or 32, because it affects the performance little. In the neural networks, we set the MLP for extracting pair relations as 2 layers with hidden dimension as 64, and the MLP for single amino acid as 2 layers with hidden dimension as 128. Following, 6 layers of transformer are stacked behind, and the final layer is the LOCS which has been discussed before.

**Figure 5:**
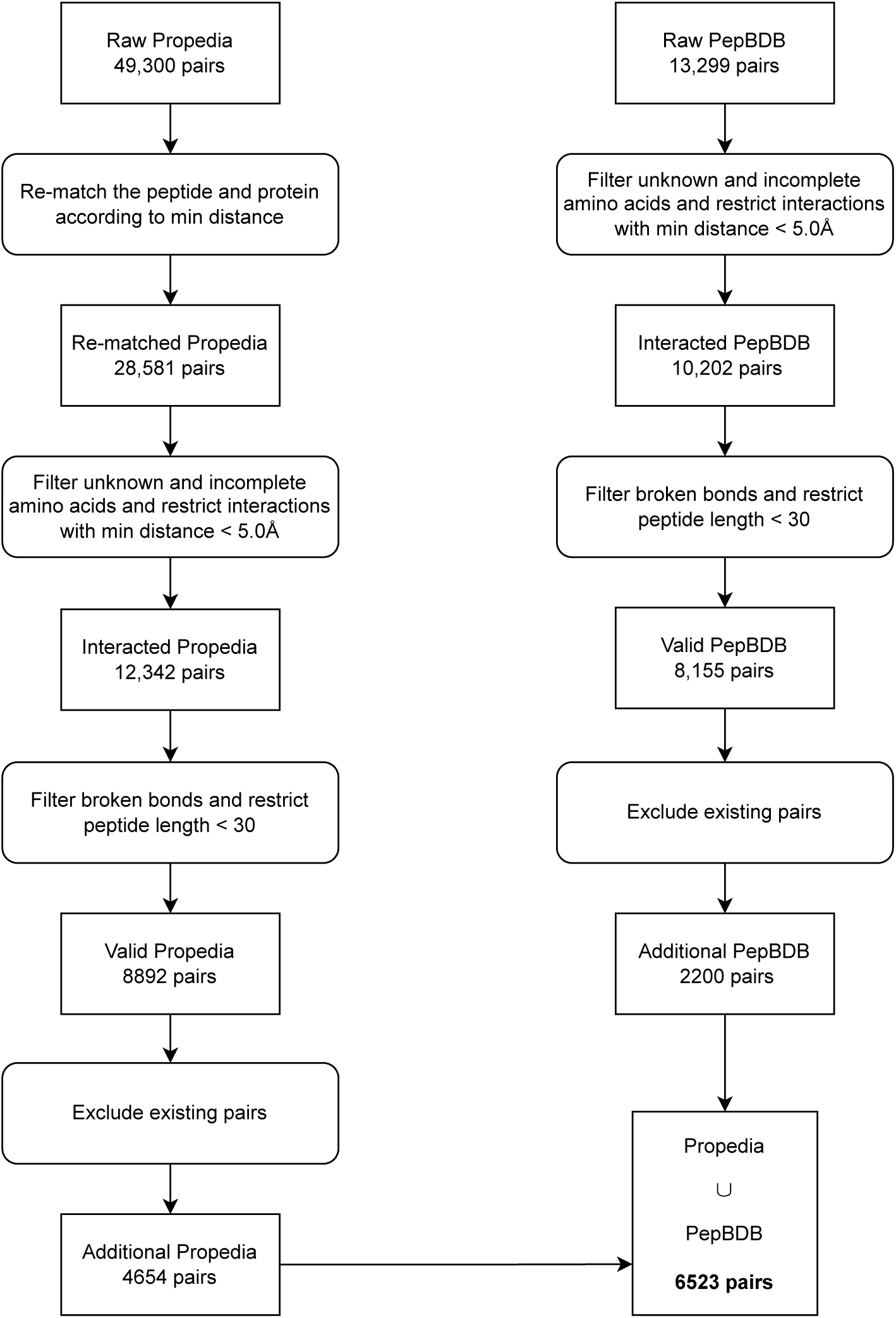
The process of screening the two datasets to expand the raw PPBench.

**Figure 6:**
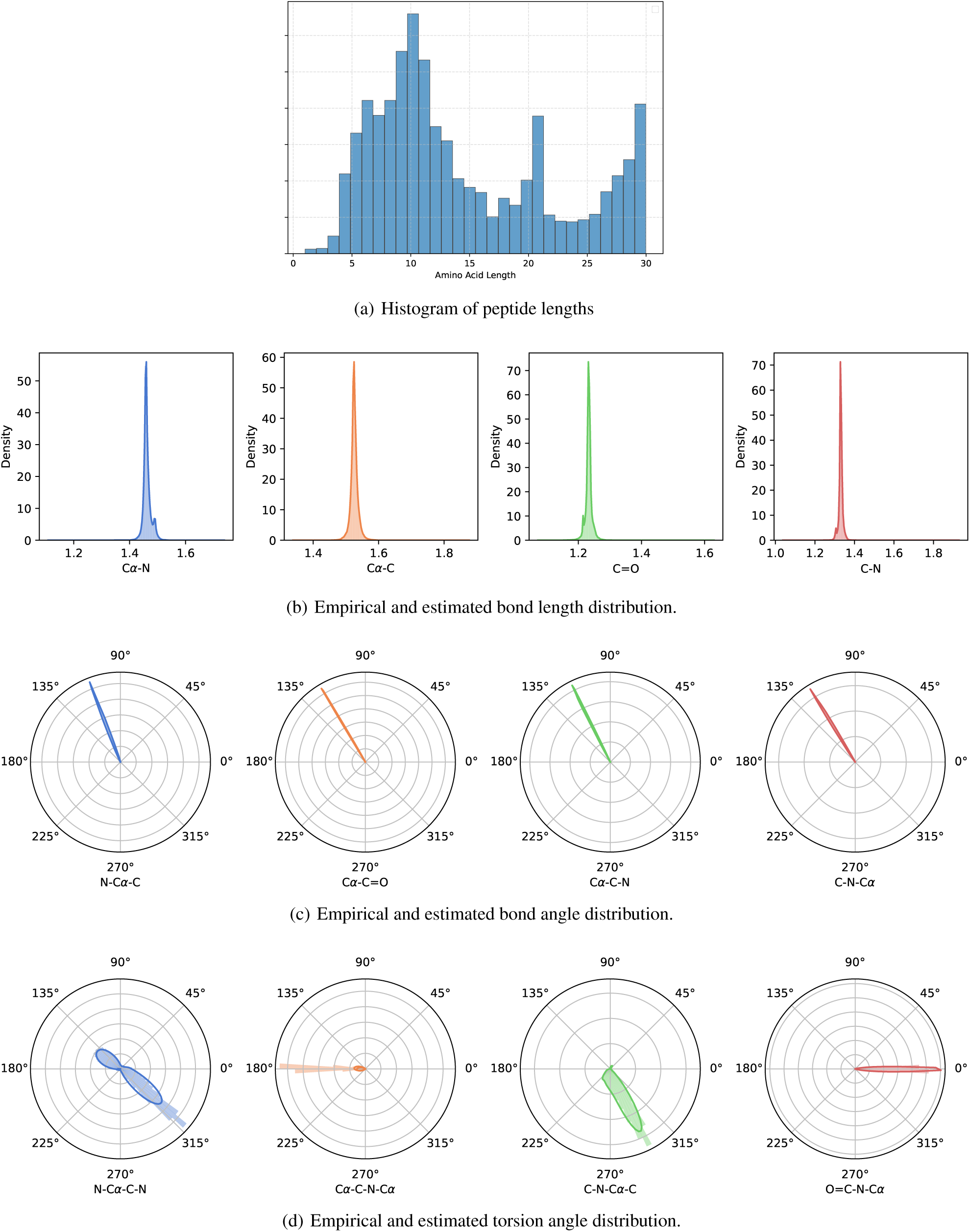
Distributions of flexible and inflexible geometries obtained by peptides in PPBench2024 datasets.

