## appendix with proof, method details and experiments for "PPFlow: Target-Aware Peptide Design with Torsional Flow Matching"

### A. Method

##### A.1. Proof of Proposition. 3.1

By Equation. 3, we can obtain $\boldsymbol{\epsilon} = \frac{\boldsymbol{\tau}_t - \boldsymbol{\mu}_t}{\sigma_t}$. Besides, from the definition of $u_t(\boldsymbol{\tau}|\boldsymbol{\tau}_1) = d\dot{\boldsymbol{\tau}}/dt$, it can be written as

$$\begin{aligned} u_t(\boldsymbol{\tau}|\boldsymbol{\tau}_1) =& \dot{\sigma}_t \boldsymbol{\epsilon} + \dot{\boldsymbol{\mu}}_t \\ =& \dot{\sigma}_t (\frac{\boldsymbol{\tau}_t - \boldsymbol{\mu}_t}{\sigma_t}) + \dot{\boldsymbol{\mu}}_t \\ =& \frac{\dot{\sigma}_t(\boldsymbol{\tau}_1)}{\sigma_t(\boldsymbol{\tau}_1)} (\boldsymbol{\tau} - \boldsymbol{\mu}_t(\boldsymbol{\tau}_1)) + \dot{\boldsymbol{\mu}}_t(\boldsymbol{\tau}_1). \end{aligned} \tag{15}$$

##### A.2. Proof of Proposition. 3.2

Firstly, we claim the disintegration of measures of $p_t(\boldsymbol{\tau})$, as $p_t(\boldsymbol{\tau}) = \prod_i p(\tau^{(i)})$. For $p_0(\boldsymbol{\tau})$, the disintegration satisfies. For $p_1(\boldsymbol{\tau})$, our parametrization assumes the torsion angles are orthogonal and the distribution is independent, so it also satisfies disintegration. In this way, it is easy to obtain an intermediate probability $p_t$ satisfies disintegration, and similar for the conditional probability $p_t(\boldsymbol{\tau}|\boldsymbol{\tau}_1)$.

Then we can factorizes the metric on $\mathbb{T}^N$ into $\mathbb{S} \times \cdots \times \mathbb{S}$, and $p(\tau) \in \mathcal{P}(\mathbb{S})$.

Let $u_t = \mathbb{E}_{\tau_0 \sim p_0, \tau_1 \sim p_1} \left[ \frac{p_t(\tau|\tau_0,\tau_1)}{p_t(\tau)} u_t(\tau|\tau_0, \tau_1) \right]$, and

$$\begin{aligned} &\nabla_\theta (\mathbb{E}_{\tau_0 \sim p_0, \tau_1 \sim p_1, \tau \sim p_t(\tau|\tau_0,\tau_1)} [\|v_t(\tau) - u_t(\tau|\tau_0, \tau_1)\|^2] - \mathbb{E}_{\tau \sim p_t} [\|v_t(\tau) - u_t(\tau)\|^2]) \\ =& - 2\nabla_\theta (\mathbb{E}_{\tau_0 \sim p_0, \tau_1 \sim p_1, \tau \sim p_t(\tau|\tau_0,\tau_1)} \langle v_t(\tau), u_t(\tau|\tau_0, \tau_1)\rangle - \mathbb{E}_{\tau \sim p_t} \langle v_t(\tau), u_t(\tau)\rangle). \end{aligned} \tag{16}$$

Then,

$$\begin{aligned} & \mathbb{E}_{\tau \sim p_t} \langle v_t(\tau), u_t(\tau)\rangle \\ =& \int \langle v_t(\tau), u_t(\tau)\rangle p_t(\tau) d\tau \\ =& \int \langle v_t(\tau), \mathbb{E}_{p_0,p_1} \left[ \frac{p_t(\tau|\tau_0,\tau_1)}{p_t(\tau)} u_t(\tau|\tau_0,\tau_1) \right] \rangle p_t(\tau) d\tau \\ =& \int \langle v_t(\tau), \int \frac{p_t(\tau|\tau_0,\tau_1)}{p_t(\tau)} u_t(\tau|\tau_0,\tau_1) p_0(\tau_0) p_1(\tau_1) d\tau_0 \tau_1 \rangle p_t(\tau) d\tau \\ =& \int\int \langle v_t(\tau), u_t(\tau|\tau_0,\tau_1)\rangle p_t(\tau|\tau_0,\tau_1) p_0(\tau_0) p_1(\tau_1) d\tau d\tau_0 d\tau_1 \\ =& \mathbb{E}_{\tau_0 \sim p_0, \tau_1 \sim p_1, \tau \sim p_t(\tau|\tau_0,\tau_1)} \langle v_t(\tau), u_t(\tau)\rangle, \end{aligned} \tag{17}$$

where in the last equality we change the order of integration. It proves Equation. 16 equals 0.

##### A.3. Proof of Proposition. 3.3

The proof is inspired by (Köhler et al., 2020; Lin et al., 2023), as follows:

**Lemma A1.** Let $\mathrm{T}_g(\cdot)$ be the operation in SE(3), if the following update function in the ODE sampler (Sec. 3.6) for the atom level's positions are defined as

$$v_t(\{X^{(i)*}\}|\mathcal{C}_t^*) = (\{X_{t+\Delta t}^{(i)*}\} - \{X_t^{(i)*}\})/\Delta t, \tag{18}$$

in which $\{X_t^{(i)*}\} = O_t \mathrm{nerf}(\boldsymbol{\tau}_t) + \mathbf{x}^{(\mathrm{C})}$. The invariance and equivariance of the following functions in the updating process are listed as

$$\begin{aligned} v\left(\{X_t^{(i)}\}|\mathrm{T}_g(\mathcal{C}^*)\right) &= \mathrm{T}_g\left(v(\{X_t^{(i)}\}|\mathcal{C}^*)\right); \\ v\left((\{s_t^{(i)}\})|\mathrm{T}_g(\mathcal{C}^*)\right) &= v\left((\{s_t^{(i)}\})|\mathcal{C}^*\right), \end{aligned} \tag{19}$$

and the prior distribution as

$$p_0(\{X_0^{(i)*}\}|\mathrm{T}_g(\mathcal{R})) = p_0(\{X_0^{(i)*}\}|\mathcal{R}) \tag{20}$$

$$p_0(\{s_0^{(i)*}\}|\mathrm{T}_g(\mathcal{R})) = p_0(\{s_0^{(i)*}\}|\mathcal{R}). \tag{21}$$

Then the distribution which the final structures and sequences are sampled from as $p\left(\mathcal{L}|\mathcal{R}\right)$ is SE(3)-equivariant.

**Proof:** In the following, we write $\boldsymbol{X} = \{X^{(i)*}\}_{i=1}^{N_{\mathrm{PP}}}$, and $\boldsymbol{s} = \{s^{(i)}\}_{i=1}^{N_{\mathrm{PP}}}$ for notation simplicity. For the defined $v_t$, we first obtain that the update process is equivariant, as

$$\begin{aligned}
&v_t(\boldsymbol{X}|\mathrm{T}_g(\mathcal{C}_t^*))\Delta t + \mathrm{T}_g(\boldsymbol{X}_t)\\
=&\mathrm{T}_g\left(v_t(\boldsymbol{X}|\mathcal{C}_t^*)\right)\Delta t + \mathrm{T}_g(\boldsymbol{X}_t)\\
=&\mathrm{T}_g\left(v_t(\boldsymbol{X}|\mathcal{C}_t^*)\Delta t + \boldsymbol{X}_t\right)\\
=&\mathrm{T}_g\left(\boldsymbol{X}_{t+\Delta t}\right)
\end{aligned} \tag{22}$$

and for each transition kernel $p(\boldsymbol{X}_{t+\Delta t}|\mathcal{C}_t^*)$, it is SE(3)-equivariant, since

$$\begin{aligned}
&p(\mathrm{T}_g(\boldsymbol{X}_{t+\Delta t})|\mathrm{T}_g(\mathcal{C}_t^*)) &(23)\\
=&p(\mathrm{T}_g(\boldsymbol{X}_t + v_t(\boldsymbol{X}|\mathrm{T}_g(\mathcal{C}_t^*))\Delta t)|\mathrm{T}_g(\mathcal{C}_t^*)) &(24)\\
=&\int p(\mathrm{T}_g(\boldsymbol{X}_{t+\Delta t}) - \mathrm{T}_g(\boldsymbol{X}_t), \mathrm{T}_g(\boldsymbol{X}_t))|\mathrm{T}_g(\mathcal{C}_t^*))d\mathrm{T}_g(\boldsymbol{X}_t) &(25)\\
=&\int p(\mathrm{T}_g(\boldsymbol{X}_{t+\Delta t}) - \mathrm{T}_g(\boldsymbol{X}_t)|\mathrm{T}_g(\mathcal{C}_t^*))p(\mathrm{T}_g(\boldsymbol{X}_t)|\mathrm{T}_g(\mathcal{C}_t^*))d\boldsymbol{X}_t &(26)\\
=&\int p(v_t(\boldsymbol{X}|\mathrm{T}_g(\mathcal{C}_t^*))\Delta t|\mathrm{T}_g(\mathcal{C}_t^*))p(\boldsymbol{X}_t|\mathcal{C}_t^*)d\boldsymbol{X}_t &(27)\\
=&\int p(v_t(\boldsymbol{X}|\mathcal{C}_t^*)\Delta t|\mathcal{C}_t^*)p(\boldsymbol{X}_t|\mathcal{C}_t^*)d\boldsymbol{X}_t &(28)\\
=&\int p(\boldsymbol{X}_{t+\Delta t} - \boldsymbol{X}_t\}|\mathcal{C}_t^*)p(\boldsymbol{X}_t|\mathcal{C}_t^*)d\boldsymbol{X}_t &(29)\\
=&p(\boldsymbol{X}_{t+\Delta t}|\mathcal{C}_t^*) &(30)
\end{aligned}$$

And it is easy to obtained that the transition kernel $p(\boldsymbol{s}_{t+\Delta t}|\mathcal{C}_t^*)$ is SE(3)-invariant, since $v_t(\boldsymbol{s}|\mathcal{C}_t^*)$ is invariant. Besides, for $p_0(\{X_0^{(i)*}\})$ and $p_0(\{s_0^{(i)*}\})$, it is a SE(3)-invariant distribution. Therefore,

$$\begin{aligned}
&p\left(\mathrm{T}_g(\boldsymbol{X}_1), \boldsymbol{s}_1|\mathrm{T}_g(\mathcal{R})\right) &(31)\\
=&\int p\left(\mathrm{T}_g(\boldsymbol{X}_0), \boldsymbol{s}_0|\mathrm{T}_g(\mathcal{R})\right)\prod_{k=1}^{K} p\left(\mathrm{T}_g(\boldsymbol{X}_{k\Delta t}), \boldsymbol{s}_{k\Delta t}|\mathrm{T}_g(\mathcal{C}^*_{(k-1)\Delta t})\right) d\mathcal{L}_{0:(K-1)\Delta t} &(32)\\
=&\int p\left(\boldsymbol{X}_0, \boldsymbol{s}_0|\mathcal{R}\right)\prod_{k=1}^{K} p\left(\mathrm{T}_g(\boldsymbol{X}_{k\Delta t})|\mathrm{T}_g(\mathcal{C}^*_{(k-1)\Delta t})\right) p\left(\boldsymbol{s}_{k\Delta t}|\mathrm{T}_g(\mathcal{C}^*_{(k-1)\Delta t})\right) d\mathcal{L}_{0:(K-1)\Delta t} &(33)\\
=&\int p\left(\boldsymbol{X}_0, \boldsymbol{s}_0|\mathcal{R}\right)\prod_{k=1}^{K} p\left(\boldsymbol{X}_{k\Delta t}|\mathcal{C}^*_{(k-1)\Delta t}\right) p\left(\boldsymbol{s}_{k\Delta t}|\mathcal{C}^*_{(k-1)\Delta t}\right) d\mathcal{L}_{0:(K-1)\Delta t} &(34)\\
=&\int p\left(\boldsymbol{X}_0, \boldsymbol{s}_0|\mathcal{R}\right)\prod_{k=1}^{K} p\left(\boldsymbol{X}_{k\Delta t}, \boldsymbol{s}_{k\Delta t}|\mathcal{C}^*_{(k-1)\Delta t}\right) d\mathcal{L}_{0:(K-1)\Delta t} &(35)\\
=&p\left(\boldsymbol{X}_1, \boldsymbol{s}_1|\mathcal{R}\right), &(36)
\end{aligned}$$

where $K$ is the step number of the ODE sampler, and $\Delta t = 1/K$. Because $p\left(\mathrm{T}_g(\boldsymbol{X}_1), \boldsymbol{s}_1|\mathrm{T}_g(\mathcal{R})\right) = p\left(\boldsymbol{X}_1, \boldsymbol{s}_1|\mathcal{R}\right)$, which is equivalent to $p\left(\mathrm{T}_g(\mathcal{L}_1^*)|\mathrm{T}_g(\mathcal{R})\right) = p\left(\mathcal{L}_1^*|\mathcal{R}\right)$, the Lemma is proved.

Then we write $\{X_t^{(i)*}\} = O_t\mathrm{nerf}(\boldsymbol{\tau}_t) + \mathrm{x}_t^{(\mathrm{C})}$, and the following proposition can give the conditions that the transition kernels of $p(O_{t+\Delta t}|\mathcal{C}_t^*)$, $p(\mathrm{x}^{\mathrm{C}}_{t+\Delta t}|\mathcal{C}_t^*)$ and $p(\boldsymbol{\tau}_{t+\Delta t}|\mathcal{C}_t^*)$ and $p(\boldsymbol{s}_{t+\Delta t}|\mathcal{C}_t^*)$ should satisfiy.

**Lemma A2.** If $p(O_0^{(\mathrm{C})})$, $p(\mathbf{x}_0^{(\mathrm{C})})$, $p(\boldsymbol{\tau}_0)$ and $p(\boldsymbol{s}_0)$ are SE(3)-invariant, and $p(O_{t+\Delta t}^{(\mathrm{C})}|\mathcal{C}_t)$ is SO(3)-equivariant and T(3)-invariant, $p(\mathbf{x}_{t+\Delta t}^{\mathrm{C}}|\mathcal{C}_t^*)$ is SE(3)-equivariant, $p(\boldsymbol{\tau}_{t+\Delta t}|\mathcal{C}_t^*)$ and $p(\boldsymbol{s}_{t+\Delta t}|\mathcal{C}_t^*)$ is SE(3)-invariant, then $p\left(\mathrm{T}_g(\mathcal{L}_1^*)|\mathrm{T}_g(\mathcal{R})\right) = p\left(\mathcal{L}_1^*|\mathcal{R}\right)$, where $\mathcal{L}_1^* = (O_1^{(\mathrm{C})}\mathrm{nerf}(\boldsymbol{\tau}_1) + \mathbf{x}_1^{(\mathrm{C})}, \boldsymbol{s}_1)$.

**Proof:** Here for $\mathrm{T}_g$, we can decompose it as $\mathrm{T}_g = \mathrm{T}_r \circ \mathrm{T}_t$, meaning the rotation and translation opeartions as $\mathrm{SE}(3) \cong \mathrm{SO}(3) + \mathrm{T}(3)$. By this mean,

$$
\begin{aligned}
& p\left(\mathrm{T}_g(\mathcal{L}_1^*)|\mathrm{T}_g(\mathcal{R})\right) && (37)\\
=& p\left(\mathrm{T}_g(O_1^{(\mathrm{C})}\mathrm{nerf}(\boldsymbol{\tau}_1) + \mathbf{x}_1^{(\mathrm{C})}), \boldsymbol{s}_1)|\mathrm{T}_g(\mathcal{R})\right) && (38)\\
=& p\left(\mathrm{T}_g(O_1^{(\mathrm{C})}\mathrm{nerf}(\boldsymbol{\tau}_1)) + \mathrm{T}_g(\mathbf{x}_1^{(\mathrm{C})})), \boldsymbol{s}_1|\mathrm{T}_g(\mathcal{R})\right) && (39)\\
=& p\left(\mathrm{T}_g(O_1^{(\mathrm{C})}\mathrm{nerf}(\boldsymbol{\tau}_1))|\mathrm{T}_g(\mathcal{R})\right) p\left(\mathrm{T}_g(\mathbf{x}_1^{(\mathrm{C})}))|\mathrm{T}_g(\mathcal{R})\right) p\left(\boldsymbol{s}_1|\mathrm{T}_g(\mathcal{R})\right) && (40)\\
=& p\left(\mathrm{T}_g(O_1^{(\mathrm{C})}\mathrm{nerf}(\boldsymbol{\tau}_1))|\mathrm{T}_g(\mathcal{R})\right) p\left(\mathrm{T}_g(\mathbf{x}_1^{(\mathrm{C})}))|\mathrm{T}_g(\mathcal{R})\right) p\left(\boldsymbol{s}_1|\mathrm{T}_g(\mathcal{R})\right). && (41)
\end{aligned}
$$

Because $\mathrm{nerf}(\boldsymbol{\tau}_1)$ is always reconstructed with unit rotation $\mathrm{diag}(1,1,1)$ and zero-mass-centered, therefore,

$$
\begin{aligned}
& p\left(\mathrm{T}_g(O_1^{(\mathrm{C})}\mathrm{nerf}(\boldsymbol{\tau}_1))|\mathrm{T}_g(\mathcal{R})\right) && (42)\\
=& p\left(\mathrm{T}_g(O_1^{(\mathrm{C})})\mathrm{nerf}(\boldsymbol{\tau}_1)|\mathrm{T}_g(\mathcal{R})\right) && (43)\\
=& p\left(\mathrm{T}_r \circ \mathrm{T}_t(O_1^{(\mathrm{C})})\mathrm{nerf}(\boldsymbol{\tau}_1)|\mathrm{T}_g(\mathcal{R})\right) && (44)\\
=& p\left(\mathrm{T}_r(O_1^{(\mathrm{C})})\mathrm{nerf}(\boldsymbol{\tau}_1)|\mathrm{T}_g(\mathcal{R})\right) && (45)\\
=& \int p\left(\mathrm{T}_r(O_0^{(\mathrm{C})})\mathrm{nerf}(\boldsymbol{\tau}_0)|\mathrm{T}_g(\mathcal{R})\right) \prod_{k=1}^{K} p\left(\mathrm{T}_r(O_{k\Delta t}^{(\mathrm{C})})\mathrm{nerf}(\boldsymbol{\tau}_{k\Delta t})|\mathrm{T}_g(\mathcal{C}_{(k-1)\Delta t}^*)\right) dM_{0:(K-1)\Delta t} && (46)\\
=& \int p\left(O_0^{(\mathrm{C})}\mathrm{nerf}(\boldsymbol{\tau}_0)|\mathrm{T}_g(\mathcal{R})\right) \prod_{k=1}^{K} p\left(O_{k\Delta t}^{(\mathrm{C})}\mathrm{nerf}(\boldsymbol{\tau}_{k\Delta t})|\mathcal{C}_{(k-1)\Delta t}^*\right) dM_{0:(K-1)\Delta t} && (47)\\
=& p\left(O_1^{(\mathrm{C})}\mathrm{nerf}(\boldsymbol{\tau}_1)|\mathcal{R}\right) && (48)
\end{aligned}
$$

where $dM_{0:(K-1)\Delta t} = dO_{0:(K-1)\Delta t}^{(\mathrm{C})} d\boldsymbol{\tau}_{0:(K-1)\Delta t}$. For $p\left(\mathrm{T}_g(\mathbf{x}_1^{(\mathrm{C})}))\right)$ and $p\left(\boldsymbol{s}_1|\mathrm{T}_g(\mathcal{R})\right)$, the equivariance and invariance are also easy to obtain. Therefore, the Lemma A2. is proved.

Finally, **PROOF OF PROPOSITION 3.3**:

For the transition kernels which are generated by the following equations as

$$
\begin{aligned}
p(\boldsymbol{\tau}_{t+\Delta t}|\mathcal{C}_t^*) &= p(\boldsymbol{\tau}_t + v_t(\boldsymbol{\tau}|\mathcal{C}_t^*)\Delta t|\mathcal{C}_t^*);\\
p(\mathbf{x}_{t+\Delta t}^{(\mathrm{C})}|\mathcal{C}_t^*) &= p(\mathbf{x}_t^{(\mathrm{C})} + v_t(\mathbf{x}^{(\mathrm{C})}|\mathcal{C}_t^*)\Delta t|\mathcal{C}_t^*);\\
p(O_{t+\Delta t}^{(\mathrm{C})}|\mathcal{C}_t^*) &= p(O_t^{(\mathrm{C})}\exp_{O_t^{(\mathrm{C})}}(v_t(\boldsymbol{\tau}|\mathcal{C}_t^*)\Delta t)|\mathcal{C}_t^*);\\
p(\boldsymbol{s}_{t+\Delta t}|\mathcal{C}_t^*) &= p(\boldsymbol{s}_t + v_t(\boldsymbol{s}|\mathcal{C}_t^*)\Delta t|\mathcal{C}_t^*),
\end{aligned} \qquad (49)
$$

if the conditions of flow-matching vector fields in **Propostion 3.3** are all satisfied, then $p(\boldsymbol{\tau}_{t+\Delta t}|\mathcal{C}_t^*)$, $p(\mathbf{x}_{t+\Delta t}^{(\mathrm{C})}|\mathcal{C}_t^*)$, $p(O_{t+\Delta t}|\mathcal{C}_t^*)$ and $p(\boldsymbol{s}_{t+\Delta t}|\mathcal{C}_t^*)$ satisfies the conditions in **Lemma A.2** resepectively, and $v(\boldsymbol{x}|\mathcal{C}_t^*)$ satisfies **Lemma A.1**. From both perspectives, the roto-translational equivariance can be proved.

We demonstarte the first proof path, to shown that the conditions of **Lemma A.2** holds.

For $\boldsymbol{\tau}$:

$$\begin{aligned} p(\mathrm{T}_g(\boldsymbol{\tau}_{t+\Delta t})|\mathrm{T}_g(\mathcal{C}_t^*)) &= p(\mathrm{T}_g(\boldsymbol{\tau}_t + v_t(\boldsymbol{\tau}|\mathcal{C}_t^*)\Delta t)|\mathrm{T}_g(\mathcal{C}_t^*)) \\ &= p(\mathrm{T}_g(\boldsymbol{\tau}_t) + \mathrm{T}_g(v_t(\boldsymbol{\tau}|\mathcal{C}_t^*)\Delta t)|\mathrm{T}_g(\mathcal{C}_t^*)) \\ &= \int p(\mathrm{T}_g(\boldsymbol{\tau}_{t+\Delta t}) - \mathrm{T}_g(\boldsymbol{\tau}_t)|\mathrm{T}_g(\mathcal{C}_t^*))p(\mathrm{T}_g(\boldsymbol{\tau}_t)|\mathrm{T}_g(\mathcal{C}_t^*))d\mathrm{T}_g(\boldsymbol{\tau}_t) \\ &= \int p(\mathrm{T}_g(v_t(\boldsymbol{\tau}|\mathcal{C}_t^*)\Delta t)|\mathrm{T}_g(\mathcal{C}_t^*))p(\mathrm{T}_g(\boldsymbol{\tau}_t)|\mathrm{T}_g(\mathcal{C}_t^*))d\mathrm{T}_g(\boldsymbol{\tau}_t) \\ &= \int p(v_t(\boldsymbol{\tau}|\mathcal{C}_t^*)\Delta t|\mathcal{C}_t^*)p(\boldsymbol{\tau}_t|\mathcal{C}_t^*)d\boldsymbol{\tau}_t \\ &= \int p(\boldsymbol{\tau}_{t+\Delta t} - \boldsymbol{\tau}_t|\mathcal{C}_t^*)p(\boldsymbol{\tau}_t|\mathcal{C}_t^*)d\boldsymbol{\tau}_t \\ &= p(\boldsymbol{\tau}_{t+\Delta t}|\mathcal{C}_t^*) \end{aligned} \tag{50}$$

Beside, for the other three variables, the equations can be deduced similarly.

##### A.4. LoCS Updating

Because the outputs should include the gradient vectors of rotation angle, global translational vector, rotation matrix, and type probability, we employ the LoCS to output the vector fields by

$$\begin{aligned} v_t(\boldsymbol{c}^{(i)}|\mathcal{C}^*) &= \mathrm{MLP}_s(\boldsymbol{h}_i); \\ v_t(\{\phi^{(i)}, \psi^{(i)}, \omega^{(i)}\}|\mathcal{C}^*) &= \mathrm{MLP}_\tau(\boldsymbol{h}_i); \\ v_t(\mathbf{x}^{(\mathrm{C})}|\mathcal{C}^*) &= \mathrm{MLP}_{\mathrm{x}}(\boldsymbol{h}_{\mathrm{G}})O_t; \\ v_t(O^{(\mathrm{C})}|\mathcal{C}^*) &= \mathrm{tran}_{O_t}(\mathrm{MLP}_O(\boldsymbol{h}_{\mathrm{G}})), \end{aligned} \tag{51}$$

in which $\boldsymbol{h}_{\mathrm{G}} = \sum_i \boldsymbol{h}_i$ is the global representation obtained by summation, $\mathrm{MLP}_s : \mathbb{R}^D \to \mathbb{R}^{20}$, $\mathrm{MLP}_\tau : \mathbb{R}^D \to \mathbb{T}^3$, $\mathrm{MLP}_{\mathrm{x}} : \mathbb{R}^D \to \mathbb{R}^3$ and $\mathrm{MLP}_{\mathrm{x}} : \mathbb{R}^D \to \mathbb{R}^3$. $\mathrm{MLP}_O$ predicts a vector in Lie group $\mathfrak{so}(3)$, and translate it to $O_t$'s tangent space by $\mathtt{tran}_{O_t}(\cdot)$. The output vector fields satisfy the equivariance and invariance conditions in Proposition. 3.3.

#### B. Dataset Statistics

##### B.1. Data preprocess

The construction of raw `PPBench2024` is given in the Sec. 5.1. For the additional complexes from `PropediaV2.3` and `PepBDB`, we give the detailed screening process in the Figure. 5. Because in PropediaV2.3, one peptide can be paired with several chains in a protein complex, another re-matching step should be conducted first.

Then, we give an empirical distribution on the peptide lengths of `PPBench2024`, in Figure. 6(a).

##### B.2. Analysis on Geometry

As the PPFLOW models the internal redundant geometry of the peptides, here we give a statistical illustration to show the flexible geometries that need to be generated. NeRF can use the following geometries in Figure. 6(b), 6(c) and 6(d) to reconstruct the full backbone structures. For these internal geometries, it can be concluded that the torsion angles of 'N-C$\alpha$-C-N' and 'C-N-C$\alpha$-C' are the two most flexible geometries. Besides, 'C$\alpha$-C-N-C$\alpha$' is theoretically inflexible since the constraints on peptide bond. However, in the observation, we find that it will deviate from the ideal value ($\pi$) a lot (about plus and minus 7°). In this way, we include it as another flexible geometries that the model needs to generate. These three torsion angles are named $\phi$, $\psi$ and $\omega$, respectively in formal definition.

##### B.3. Experiment

Here we give the hyper-parameters and other training details. The learning rate $lr$ is $5e-5$. In all training, the max training iteration is 200000. `LambdaLR` schedule is used, with `lr_lambda` is set as $0.95 \times lr$. The batch size is set 16 or 32, because it affects the performance little. In the neural networks, we set the MLP for extracting pair relations as 2 layers with

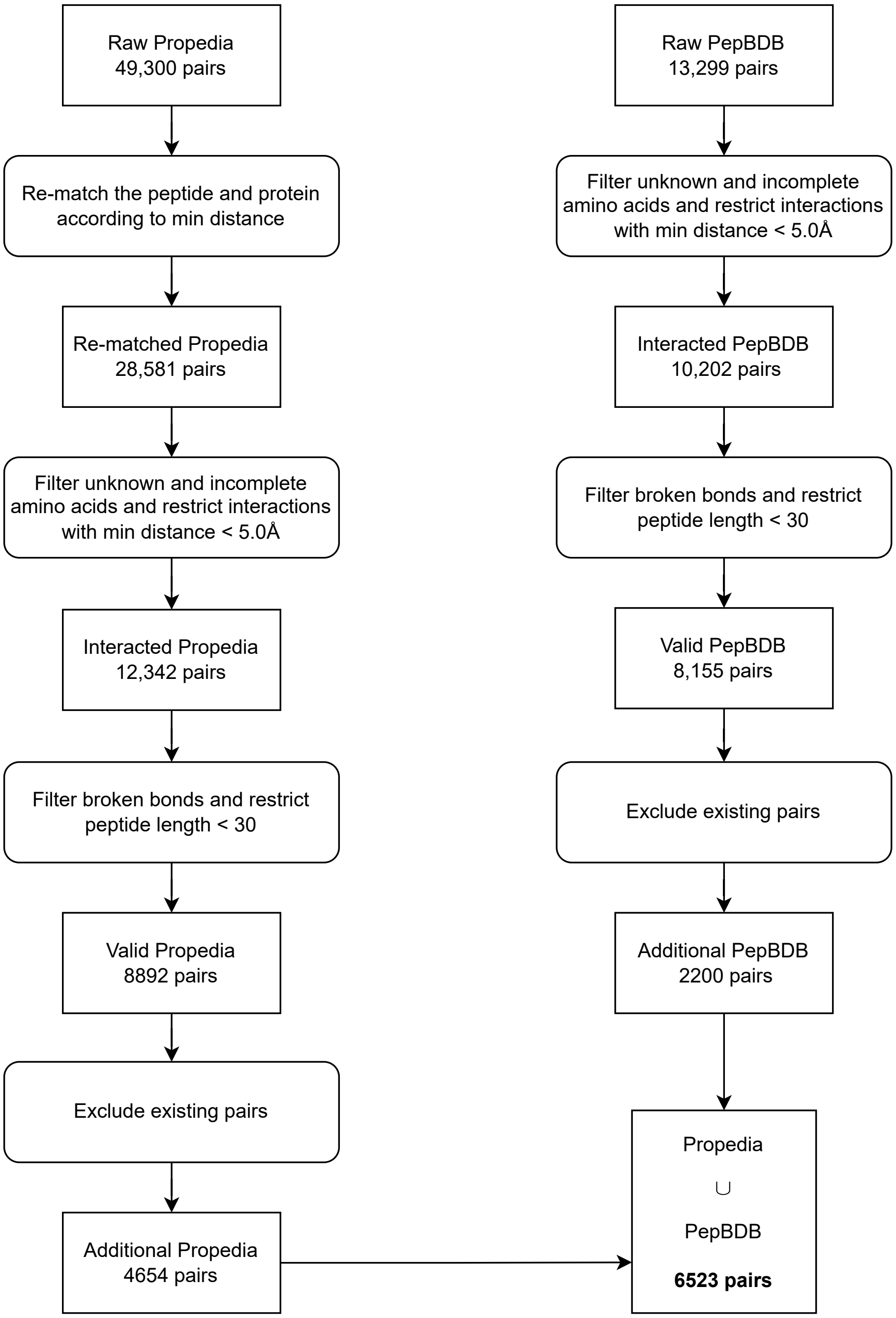


Figure 5: The process of screening the two datasets to expand the raw PPBench.

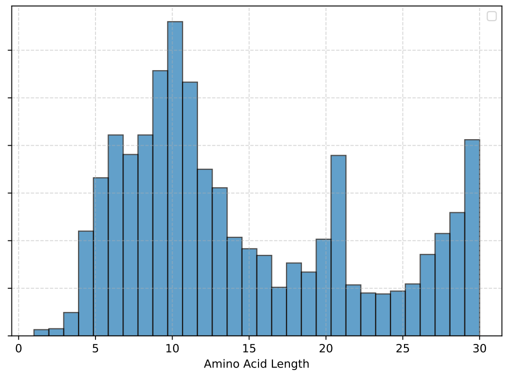


(a) Histogram of peptide lengths

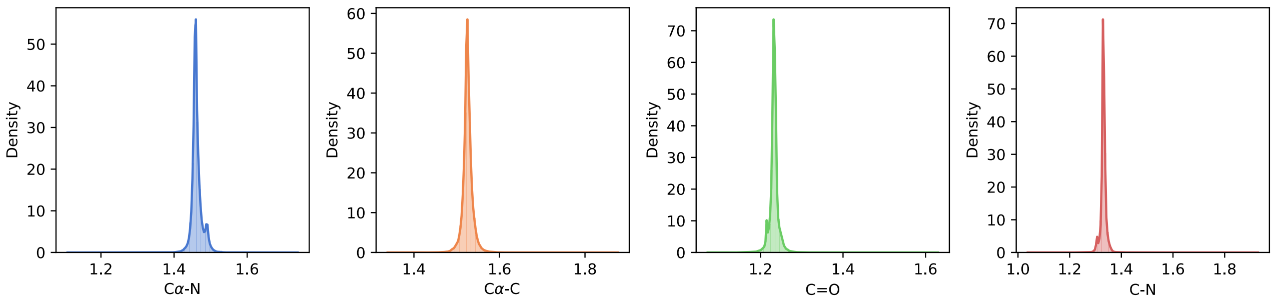


(b) Empirical and estimated bond length distribution.

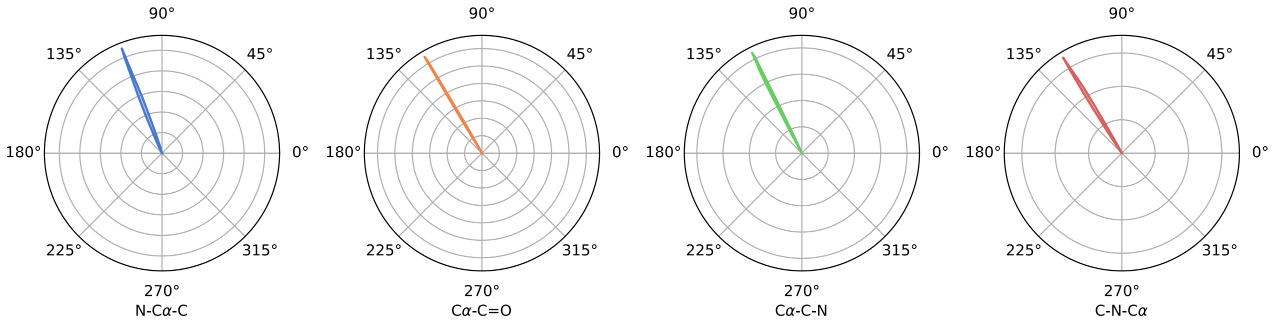


(c) Empirical and estimated bond angle distribution.

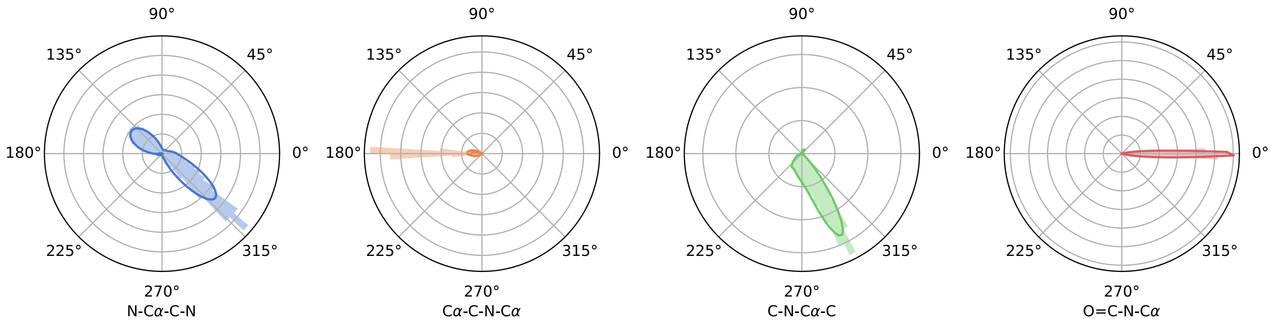


(d) Empirical and estimated torsion angle distribution.

Figure 6: Distributions of flexible and inflexible geometries obtained by peptides in `PPBench2024` datasets.

hidden dimension as 64, and the MLP for single amino acid as 2 layers with hidden dimension as 128. Following, 6 layers of transformer are stacked behind, and the final layer is the LOCS which has been discussed before.
