## Supplementary figures and images for "PPFlow: Target-Aware Peptide Design with Torsional Flow Matching"

### ppflow workflow

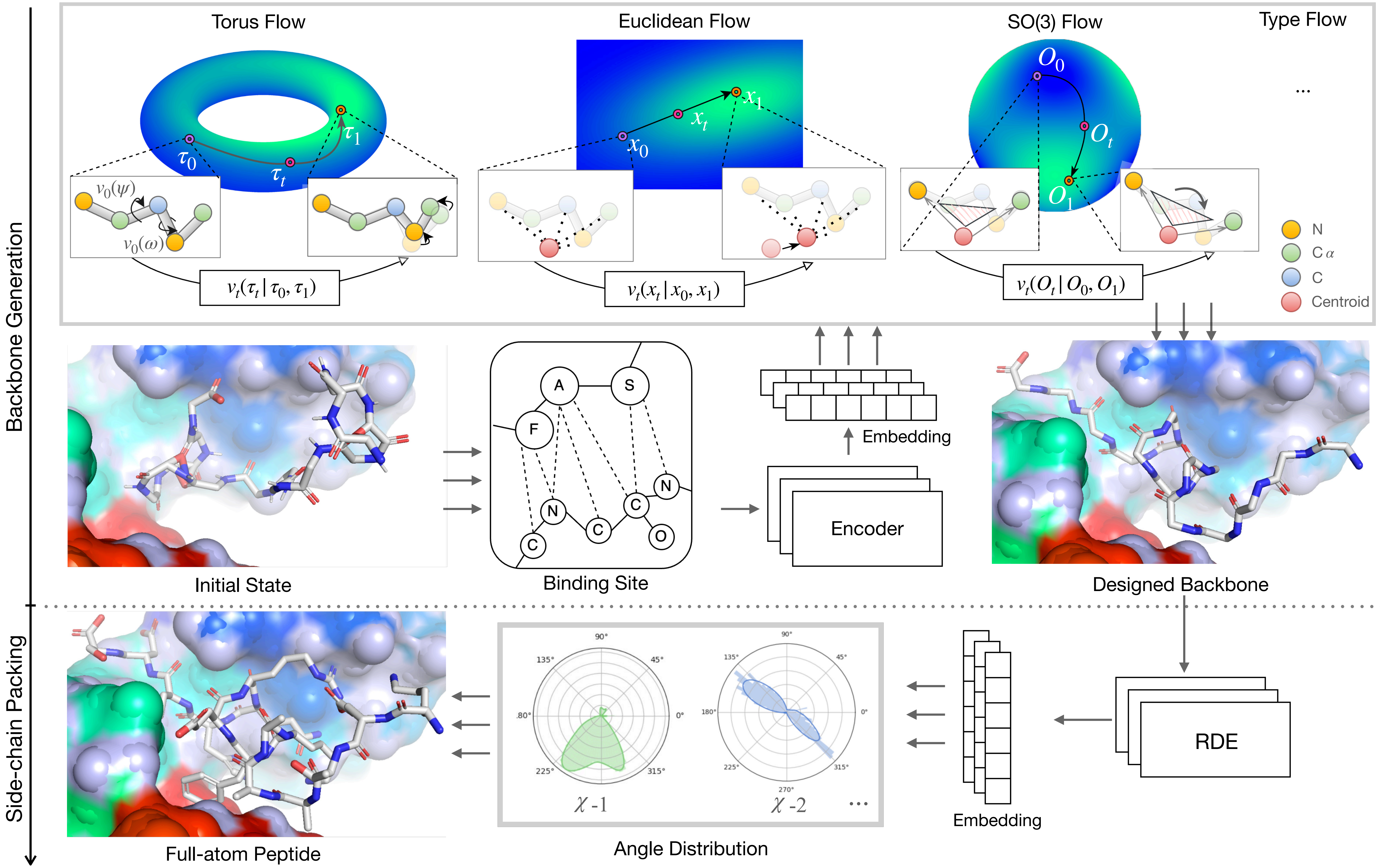
